# Endomitosis controls tissue-specific gene expression during development

**DOI:** 10.1101/2021.03.12.435137

**Authors:** Lotte M van Rijnberk, Reinier L van der Palen, Erik S Schild, Hendrik C Korswagen, Matilde Galli

## Abstract

Polyploid cells contain more than two copies of the genome and are found in many plant and animal tissues. Different types of polyploidy exist, in which the genome is confined to either one nucleus (mononucleation) or two or more nuclei (multinucleation). Despite the widespread occurrence of polyploidy, the functional significance of different types of polyploidy are largely unknown. Here, we assess the function of multinucleation in *C. elegans* intestinal cells through specific inhibition of binucleation without altering genome ploidy. Through single worm RNA sequencing, we find that binucleation is important for tissuespecific gene expression, most prominently for genes that show a rapid upregulation at the transition from larval development to adulthood. Regulated genes include vitellogenins, which encode yolk proteins that facilitate nutrient transport to the germline. We find that reduced expression of vitellogenins in mononucleated intestinal cells leads to progeny with developmental delays and reduced fitness. Together, our results show that binucleation facilitates rapid upregulation of intestine-specific gene expression during development, independently of genome ploidy, underscoring the importance of spatial genome organization for polyploid cell function.

## Introduction

Polyploidization occurs in many plant and animal cells as part of a developmental program, where it is crucial to increase cell size and metabolic output, as well as to maintain the barrier function of certain tissues. For example, polyploidization of the giant neurons of *Limax* slugs allows them to reach the enormous size they need to transmit signals over large distances and polyploidization of the *Drosophila* subperineural glial cells ensures the integrity of the blood-brain barrier during organ growth [1,2]. In mammals, polyploid cells are present in many organs, such as the liver, blood, skin, pancreas, placenta and mammary glands, where they play important functions in tissue homeostasis, regeneration and in response to damage [3,4]. Interestingly, polyploid cells can either be mononucleated, such as megakaryocytes and trophoblast giant cells [5,6], or they can be multinucleated, such as cardiomyocytes and mammary epithelial cells [7,8], but how genome partitioning in either one or multiple nuclei affects polyploid cell function is yet unknown.

Somatic polyploidy can arise either by cell fusion or by non-canonical cell cycles in which cells replicate their DNA but do not divide. Two types of non-canonical cell cycles that result in polyploidy have been described: endoreplication and endomitosis [9]. In endoreplication, M phase is skipped, resulting in cycles of DNA replication (S phase) and gap (G) phases without intervening mitosis and cytokinesis. Endoreplicative cell cycles result in large, mononucleated cells. In endomitosis, cells do enter mitosis, but do not undergo cell division, resulting in polyploid cells with either a single nucleus or two nuclei, depending on whether M phase is aborted before or after initiation of sister chromosome segregation (which normally occurs during anaphase) [3,4,10]. The existence of multiple types of polyploid cells (e.g. mononucleated or binucleated), suggests that there may be a functional difference between these different types of polyploidy. However, testing the functional significance of multinucleation has been challenging due to the complexity of many polyploid tissues and a lack of tools to specifically alter non-canonical cells cycles without affecting any other cells or tissues.

The *C. elegans* intestine provides an ideal model system to study distinct types of polyploidy, as intestinal cells undergo both endomitosis and endoreplication cycles in a highly tractable manner at defined moments during larval development [11]. Such consecutive cycles of endomitosis and endoreplication give rise to large, binucleated polyploid cells that make up the adult intestine. Here, we develop a method to inhibit intestinal endomitosis using auxin-inducible degradation of key mitotic regulators, allowing us to study the function of binucleation at both the cellular and tissue level. We find that animals with mononucleated instead of binucleated intestinal cells have decreased fitness due to defects in the expression of a group of tissue-specific genes, including the vitellogenin genes. Vitellogenin genes are rapidly expressed during the maturation of intestinal cells at the end of larval development, indicating that rapid upregulation of tissue-specific genes requires binucleation of polyploid cells. Importantly, this rapid upregulation of intestine-specific genes is important to support progeny development. Together, our results show that binucleation is important for correct functioning of a polyploid tissue, and that partitioning of genomes into multiple nuclei allows efficient and rapid upregulation of gene expression during development.

## Results

### Auxin-inducible degradation of mitotic regulators prevents binucleation

To investigate the function of binucleation, we developed a system in which we can specifically perturb intestinal endomitosis, without affecting other cell cycles within developing *C. elegans* larvae. We employed two mechanistically distinct approaches to block binucleation of the intestine; we either depleted CDK-1, which is essential for mitotic entry, or KNL-1, a conserved kinetochore protein that is required for chromosome segregation during M phase. CDK-1 inhibition is known to be a key mechanism to initiate endoreplication, and degrading CDK-1 essentially converts the endomitosis cycle to an endoreplication cycle, preventing mitotic entry and subsequent binucleation [5,9,12,13]. In contrast, KNL-1 inhibition does not affect mitotic entry but prevents chromosome segregation and partitioning of sister chromatids into two nuclei [14,15]. To deplete CDK-1 and KNL-1 specifically in intestinal cells and only during the time when endomitosis occurs, we made use of the auxin-inducible degradation system (Figure 1A). In this system, proteins tagged with an auxin-inducible degron (AID) are degraded only in the presence of auxin and the E3 ligase TIR1 [16–18]. By using a strain that expresses TIR1 exclusively in the intestine and exposing animals to auxin only during the time that endomitosis occurs, we can specifically inhibit intestinal binucleation without affecting canonical cell cycles of intestinal or other cells.

**Figure 1.**
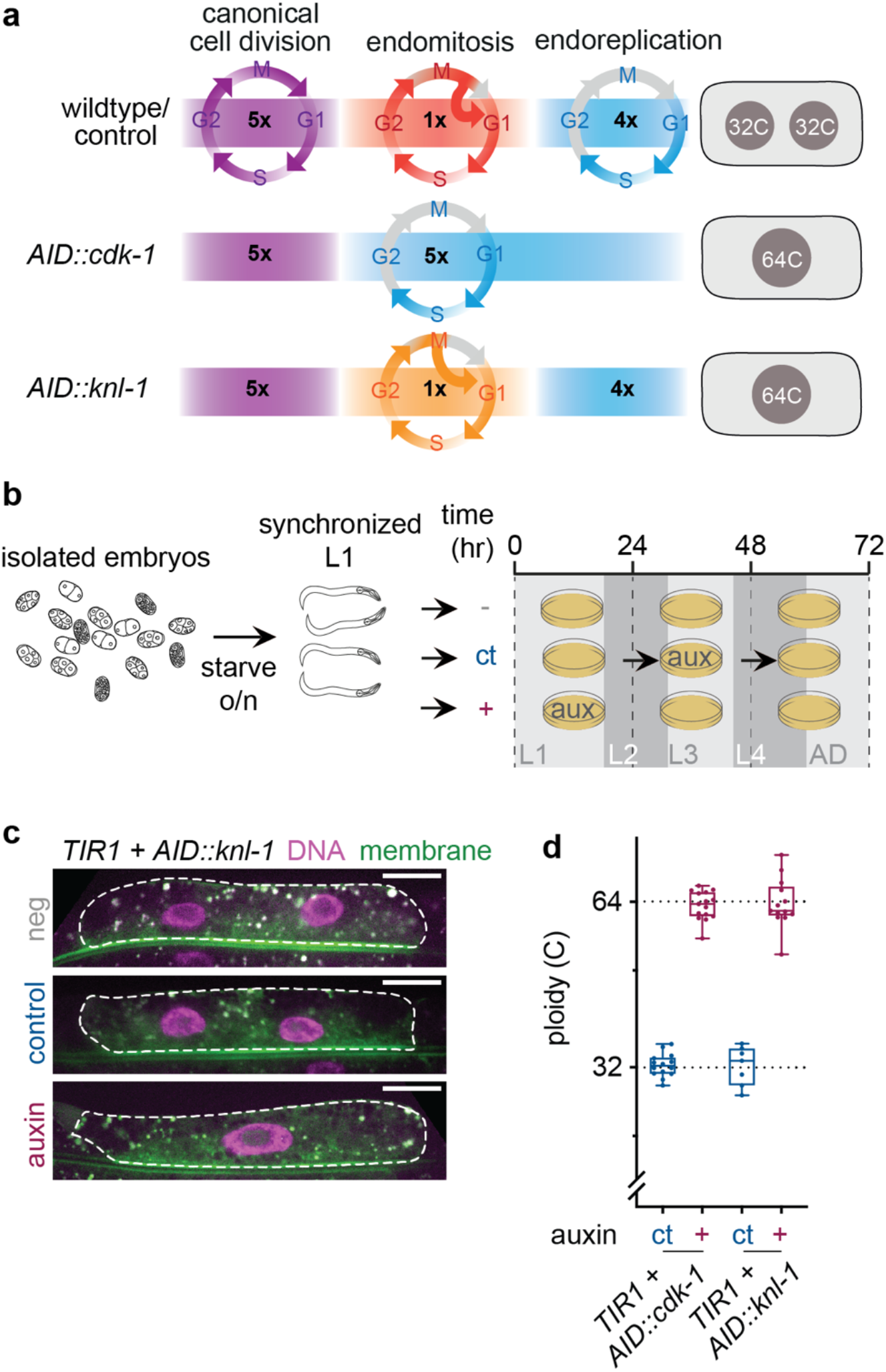
Inducible and tissue-specific degradation of mitotic proteins specifically prevents binucleation of the C. elegans intestine. **a.** Overview of intestinal cell cycles. During embryogenesis, 20 intestinal cells are formed by canonical cell cycles. After hatching, intestinal cells undergo one round of endomitosis creating a binucleated cell, and several rounds of endoreplication that increase the ploidy of these nuclei. Upon inhibition of either CDK-1 or KNL-1 during endomitosis using the auxin-inducible degradation system, endomitotic binucleation is blocked and a mononucleated polyploid cell is generated. Importantly, cellular ploidy remains the same as wildtype conditions. **b.** Schematic overview of experimental procedure. Mixed stage embryos are isolated from gravid hermaphrodites containing intestinal expressed TIR1 and either *AID::cdk-1* or *AID::knl-1* and starved overnight. The population of starved L1 stage animals is split into three conditions and grown without auxin (-), under control (ct) or auxin (+) conditions. **c.** Fluorescent images of intestinal H2B-mCherry (DNA) and GFP-PH (membrane) in intestinal ring 3 of worms that were never grown on auxin (neg); grown on auxin during later larval development (control) or grown on auxin at the moment of endomitosis (auxin). Dashed line indicates cell outline. Scale bar is 20 μm. **d.** Quantification of intestinal nuclear ploidy in binucleated control (ct, *n* = 49 for *AID::CDK-1* strain and n = 28 for *AID::KNL-1* strain) or mononucleated intestinal cells (+ auxin, *n* = 29 for *AID::CDK-1* strain and n = 26 cells for *AID::KNL-1* strain). Ploidy is measured by total fluorescent intensity of propidium iodide DNA staining in intestinal ring 3 nuclei (Int3D and Int3V), and normalized to proximal 2C nuclei. Each dot represents the average ploidy of individual Int3D/V nuclei in one animal. Boxplots indicate the median and 25^th^-75^th^ percentile, error bars indicate min to max values and individual values are shown as dots.

We generated AID knock-ins on the *cdk-1* and *knl-1* genes using CRISPR-mediated gene targeting and tested whether auxin induced depletion of CDK-1 or KNL-1 during endomitosis was able to block binucleation. In wildtype animals, endomitosis takes place at the end of the first larval stage in 12-14 of the 20 intestinal cells, resulting in 20 intestinal cells with 32-34 nuclei. To control for non-specific effects of auxin on development, we included an auxin control in each experiment, in which animals were treated with auxin for the same duration, but during a time in development in which CDK-1 or KNL-1 are not required. To assess the effect of auxin treatment on intestinal binucleation, we analyzed animals grown without auxin (hereafter referred to as untreated animals or “-”), with auxin during endomitosis (hereafter referred to as auxin-treated animals or ”+”), or in the auxin-control condition (hereafter referred to as auxin-control animals or “ct”) and found that auxin-treated animals contain 20 intestinal nuclei, in contrast to the 32-34 nuclei that are present in untreated and auxin-control animals (Figure 1B-C and data not shown). We next performed a quantitative DNA staining using propidium iodide (PI) on *AID::cdk-1* and *AID::knl-1* animals and found that auxin-treated *AID::cdk-1* and *AID::knl-1* animals have a 64C DNA content, which is exactly double the nuclear ploidy of controls (Figure 1D). Thus, by inhibiting CDK-1 and KNL-1 during endomitosis, we can block binucleation and generate animals with mononucleated intestinal cells with the same cellular ploidy as wildtype binucleated intestinal cells.

Depletion of KNL-1 is known to prevent chromosome segregation in anaphase by blocking formation of kinetochore microtubule attachments [14]. To exclude that KNL-1 knockdown results in formation of micronuclei caused by chromosome missegregations, which can lead to DNA damage, cellular stress and potentially arrest cells in the following cell cycle [19–22], we performed live-cell imaging of fluorescently-labeled chromosomes in KNL-1 depleted cells. In all cells, KNL-1 depletion prevented chromosome segregation and resulted in mononucleation, but did not extend the duration of mitosis or result in the formation of micronuclei (Supplemental Figure 1A-B, Supplemental Movie 1,2). After endomitosis, cells go through multiple rounds of endoreplication in which newly formed replicated chromosomes remain clustered together. Wildtype intestinal cells that have undergone endomitosis contain two nuclei, which each contain two sets of chromosome clusters, which can be visualized by chromosomal LacO/LacI tagging (Supplemental Figure 1C). As expected, we observed on average four chromosome clusters in nuclei of mononucleated KNL-1 depleted cells (Supplemental Figure 1D), indicating that all copies of the labeled chromosome were present in a single nucleus. Finally, using a fluorescent cell-cycle marker (*Pges-1:: CYB-1^DB^::mCherry*), we found that preventing binucleation did not affect the timing of subsequent endoreplicative cell cycles (Supplemental Figure 1E). Taken together, our results indicate that KNL-1 depletion prevents binucleation without inducing detectable chromosomal or cell cycle aberrancies. Thus, this system provides a unique opportunity to study the function of binucleation, without altering the ploidy or number of cells in the tissue of interest.

### Intestinal mononucleation decreases the nuclear surface-to-volume ratio, but does not affect cell size or morphology

To investigate whether mononucleation influences intestinal cell size or morphology, we used fluorescent markers in the *AID::knl-1* strain to visualize the cell membranes and intestinal lumen. We found no effect of intestinal mononucleation on animal size, cell size or lumen morphology (Figure 2A-F). Because polyploidization has also been shown to influence nuclear morphology [23], we investigated the effect of binucleation on nuclear geometry using a fluorescently-labeled nuclear pore protein (*NPP-9::mCherry*) as a marker for the nuclear membrane (Figure 3G). We measured nuclear size and found that the nuclear volume was increased with a factor of ~2.5 in mononucleated cells, indicating that per cell, the total nuclear volume has more than doubled (Figure 2I). The nuclear surface area was also increased in mononucleated cells, but to a lesser extent, with a factor of two (Figure 2J). As a consequence, the surface-to-volume ratio decreased, indicating that less surface area is available per volume in mononucleated cells (Figure 2K). To test whether this effect is compensated to any extent by shape alterations or membrane invaginations that enlarge nuclear surface area, such has been observed in other polyploid cells [23], we also measured the ratio between the circumference and area of nuclear sections, but observed a similar effect (Supplemental Figure 2A-D), indicating that there are no large invaginations that compensate for the amount of available nuclear surface area per volume. Thus, although mononucleation does not affect intestinal cell size or morphology, it produces nuclei that are more than twice as big and have an altered surface-to-volume ratio, which could potentially influence nuclear functions.

**Figure 2.**
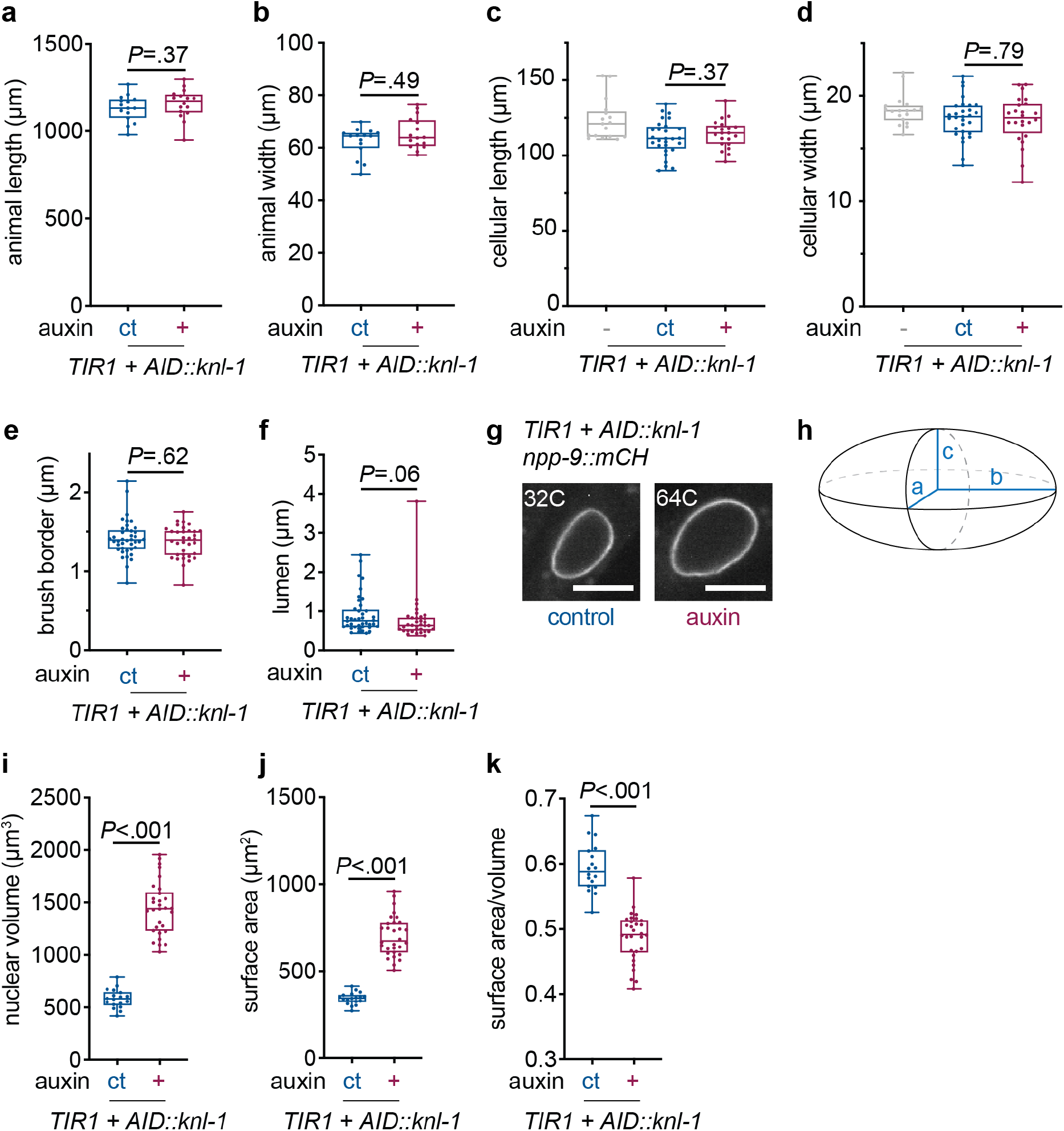
Intestinal cell mononucleation does not influence worm size, cell size or intestinal morphology, but generates nuclei that are more than twice as big. **a-b.** Total body length **(a)** and width **(b)** for young adult worms carrying an intestinally expressed TIR1, and *AID::knl-1*, grown under control conditions (ct, *n* = 15) or in the presence (+, *n* = 16) of auxin. Boxplots indicate the median and 25^th^-75^th^ percentile, error bars indicate min to max values and individual values are shown as dots. *P* values were calculated by standard unpaired student’s t-test and Mann-Whitney test, respectively. **c-d.** Cellular length **(c)** and width **(d)** of Int3D/V cells of young adult worms grown without auxin (-, *n* = 36), under control conditions (ct, *n* = 58) or in the presence of auxin (+, *n* = 24). Boxplots indicate the median and 25^th^-75^th^ percentile, error bars indicate min to max values and individual values are shown as dots. *P* values were calculated by standard unpaired student’s t-test. **e-f.** Intestinal brush border **(e)** and lumen **(f)** width in intestinal ring 3 of young adult worms grown under control (ct, *n* = 40) or auxin (+, *n* = 34) conditions. Boxplots indicate the median and 25^th^-75^th^ percentile, error bars indicate min to max values and individual values are shown as dots. *P* values were calculated by Mann-Whitney test. **g.** Fluorescent images of intestinal NPP-9::mCherry localization in the nuclear membranes of Int3 cells of adult worms that were grown under control or auxin conditions, corresponding to a 32C and 64C DNA content, respectively. Scale bar represents 10 μm. **h.** Schematic depicting an ellipsoid shape and radii a, b and c used to calculate nuclear volume and surface area. **i-k.** Boxplots depicting the average nuclear volume **(i)**, surface area **(j)** and surface-area-to-volume ratio **(k)** in binucleated (ct, *n* = 60 cells, 18 animals) or mononucleated (+, *n* = 56 cells, 30 animals) cells, per worm, as calculated with parameters mentioned in **(h)**. Boxplots indicate the median and 25^th^-75^th^ percentile, error bars indicate min to max values and individual values are shown as dots. *P* values were calculated by Mann-Whitney **(i-j)** or unpaired student’s t-test **(k)**.

**Figure 3.**
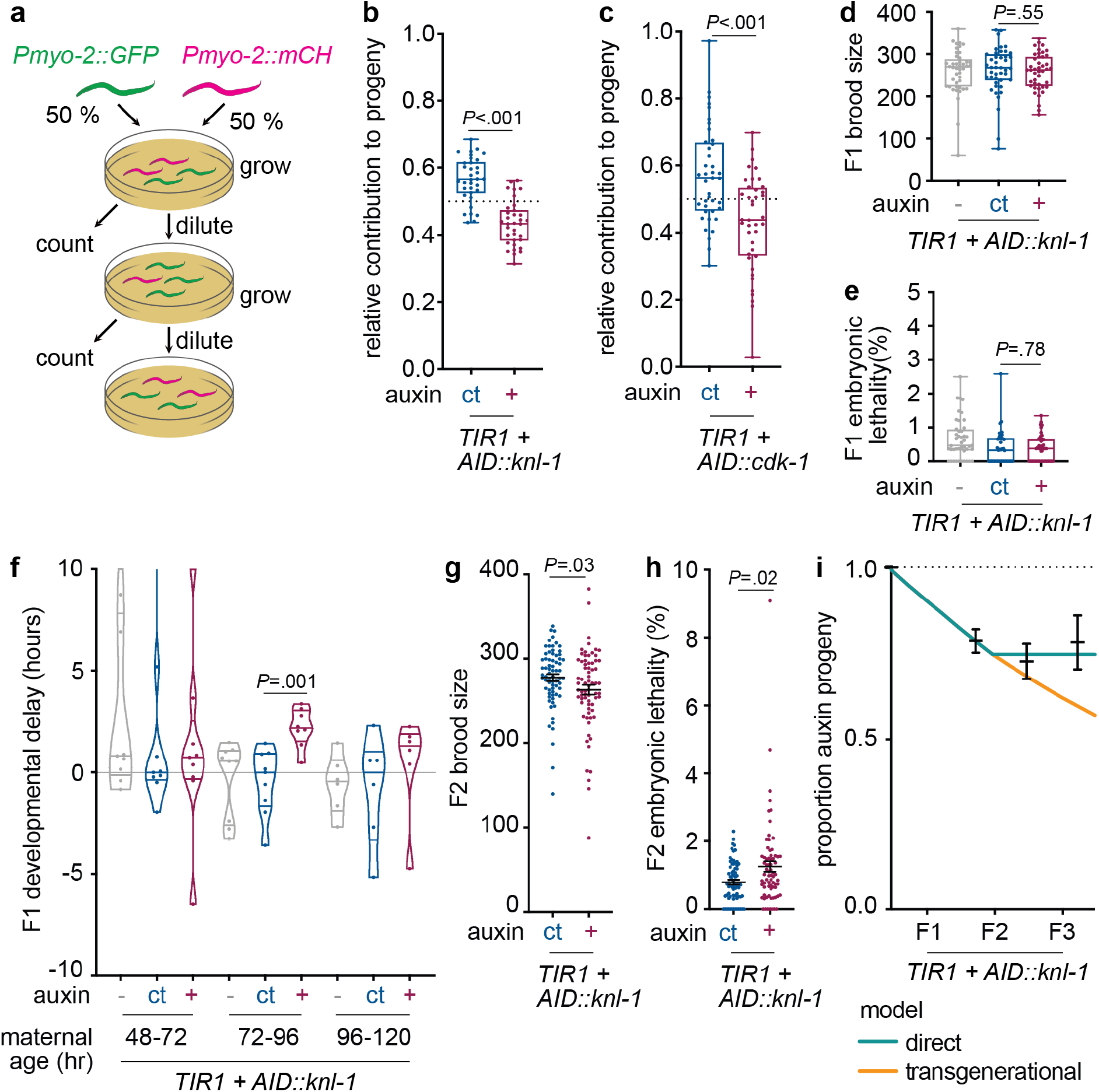
Inhibition of intestinal binucleation reduces relative reproductive fitness. **a.** Overview of the relative fitness assay. L4 animals grown under auxin or control conditions carrying a pharyngeal marker in either red (*Pmyo-2::mCherry*) or green (*Pmyo-2::GFP*) are transferred in equal amounts to a single plate. Replicate experiments include worms from the auxin and control conditions labelled in either condition. Worms are then grown until the plate is full or starved, after which the worms are chunked to a new plate, diluting the population, after which the proportion of *mCherry+* and *GFP+* progeny is counted. **b-c.** Boxplots showing the normalized proportion of progeny originating from worms with a wildtype binucleated (ct, *n* = 40 plates/280 worms and 34 plates/238 worms) or mononucleated (+, *n* = 40 plates/280 worms and 34 plates/238 worms) intestine in relative fitness assays using either the *AID::cdk-1* **(b)** or *AID::knl-1* alleles **(c)** to block binucleation, in two replicate experiments. Boxplots indicate the median and 25^th^-75^th^ percentile, error bars indicate min to max values and individual values are shown as dots. **d-e.** Boxplots showing total brood size **(d)** and embryonic lethality **(e)** of worms carrying an intestinally expressed TIR1 and AID::KNL-1 that were grown in the absence of auxin (-, *n* = 44 plates), under control conditions (ct, *n* = 43 plates) or in the presence of auxin (+, *n* = 43 plates), in three replicate experiments. Amounts of eggs and hatched worms were counted for three days of egg-laying. Boxplots indicate the median and 25^th^-75^th^ percentile, error bars indicate min to max values and individual values are shown as dots. **f.** Violin boxplots depicting progeny growth rates from worms with increasing maternal age, grown without auxin (-, *n* = 9 plates), under control conditions (ct, *n* = 9 plates) or in the presence of auxin (+, *n* = 9 plates), in three replicate experiments. Horizontal lines indicate the median and 25^th^-75^th^ percentile, violin plots extend to min and max values and individual values are shown as dots. **g-h.** Scatter plots of brood size and embryonic lethality in the second generation of worms (F2) that originated from animals grown under control (ct, *n* = 70 plates) or auxin (+, *n* = 71 plates) conditions. Amounts of eggs and hatched worms were counted for three days of egg-laying. Line and error bars represent mean and SEM. **i.** Exponential growth model assuming a direct (blue) or transgenerational (orange) effect of blocking binucleation on reproductive fitness. Experimental data (mean ± s.e.m.) of the relative fitness assay over three generations is depicted in black. **(b-h)** *P* values were calculated by Mann-Whitney test.

### Intestinal binucleation enhances C. elegans fitness

To assess whether intestinal binucleation is functionally important for *C. elegans*, we first performed a competitive fitness assay with animals that have either mononucleated or binucleated intestinal cells. To this end, we generated strains with either a *Pmyo-2::mCherry* or a *Pmyo-2::GFP* pharyngeal marker in addition to the *AID::knl-1* or *AID::cdk-1* alleles. This allowed us to follow the progeny of animals with mononucleated or binucleated cells over several generations. By mixing equal amounts of animals with mononucleated or binucleated intestines on plates and counting the proportion of GFP and mCherry positive progeny after several generations, we could determine how binucleation influences reproductive fitness (Figure 3A). It is important to note that auxin was only added during development of the parental worms, and that progeny were not treated with auxin and their intestinal cells were thus binucleated. To control for fitness differences due to the *Pmyo-2::mCherry* or *Pmyo-2::GFP* markers, we included experiments with reversed fluorescent markers and used measurements of the fitness difference between *Pmyo-2::mCherry* and *Pmyo-2::GFP* control worms to normalize our data (see Online Methods for details). Using the *AID::knl-1* strain, we found that after one generation, animals with a wildtype intestine had given rise to an average of 57.0% (±14.4%) of the population, while animals with a mononucleated intestine only gave rise to 43.0% (±14.4%) of the population (Figure 3B). Similar numbers were obtained with the *AID::cdk-1* strain (Figure 3C). These results demonstrate that animals with binucleated intestines have a significant fitness advantage over animals with mononucleated intestines.

To understand how perturbation of binucleation affects fitness, we investigated several aspects of worm growth and reproduction upon inhibition of binucleation. When analyzing *AID::knl-1* and *AID::cdk-1* animals, we noticed that the *AID::cdk-1* strain had smaller brood sizes and increased embryonic lethality compared to wildtype animals, both in the presence or absence of auxin, suggesting that the AID tag compromises CDK-1 function resulting in a weak hypomorph. Nonetheless, when comparing animals with mononucleated or binucleated intestines in either the *AID::cdk-1* or *AID::knl-1* background, we found no effect of mononucleation on brood size and embryonic lethality (Figure 3D-E, Supplemental Figure 3A). However, we did find a significant delay in progeny growth when examining eggs that were laid between 72 and 96 hours of development, corresponding to the first and second day of adulthood (Figure 3F). This delay was not due to growing animals on auxin or the presence of Tir1, as worms lacking the *AID::knl-1* or *AID::cdk-1* alleles did not show developmental delays or reproductive defects (Supplemental Figure 3B-D). Strikingly, animals derived from grandmothers with mononucleated intestines showed a decrease in brood size and increased embryonic lethality compared to controls (Figure 3G-H). To test whether these developmental effects could account for the observed decreases in fitness, we used an exponential growth model to predict the relative fitness of worms with a mononucleated intestine based on our growth and reproduction measurements. In this model, we differentiated the possibilities of an intergenerational effect of blocking binucleation in the intestine, where only the first generation of progeny is affected, and a transgenerational effect that lasts for several generations. Comparing this model with our experimental data revealed two things. First, the plateau in the proportion of progeny coming from mothers with a mononucleated intestine indicates that there is an intergenerational rather than a transgenerational effect on progeny fitness (Figure 3I). Secondly, the fitness decrease that we measured closely matches the model, suggesting that decreases in progeny growth and reproduction fully explain the difference in relative fitness. Taken together, our data shows that intestinal binucleation is important for reproductive fitness and that blocking binucleation decreases progeny growth and reproduction.

### Binucleation is important for adult intestinal transcription

We hypothesized that the observed fitness phenotype may be caused by transcriptional changes of the intestine caused by an alteration of the nuclear size or surface-to-volume ratio. To investigate possible transcriptional differences between young adults with mononucleated or binucleated intestines, we performed single-worm RNA sequencing of *AID::kln-1* and *AID::cdk-1* auxin-treated and auxin-control animals. Since the intestine is one of the most transcriptionally active and largest tissues in the worm, making up roughly one-third of the animals volume [24], we anticipated that changes in transcription in the intestine would be detectable in whole worm sequencing. We performed differential gene expression analysis for 68 (*AID::knl-1*) and 75 (*AID::cdk-1*) young adult (72 hr) auxin-treated or auxin-control animals (see Online Methods). For the *AID::knl-1* animals alone, 16 percent of genes included in the analysis showed significant and substantial differential expression between worms with a mononucleated or binucleated intestine (420/2638 genes, Figure 4A, Supplemental Figure 4A). Sequencing of *AID::cdk-1* animals revealed fewer differentially expressed genes between animals with mononucleated or binucleated intestines, which is likely due to a lower complexity of these samples (Supplemental Figure 4B-E). To identify differentially expressed genes that could explain the fitness differences between animals with binucleated or mononucleated intestines, we focused on the overlap in differential gene expression between the *AID::knl-1* and *AID::cdk-1* strains (Supplemental Figure 4F). We found a strong enrichment (*P*=2e-7) in the overlap between genes significantly and substantially downregulated in both strains, consisting of 15 genes (Figure 4B-C). One of the genes that stood out was *vit-2*, one of the six *C. elegans* vitellogenin genes whose levels have been shown to correlate strongly with the growth and fitness traits of *C. elegans* progeny [25]. Vitellogenins are highly expressed in adult intestines and are essential to mobilize lipids in the intestine for transport to the developing oocytes in the germline, where they contribute to progeny development [26–29]. Downregulation of one or multiple of these vitellogenins could therefore explain the phenotypes that we observed in the progeny of animals with mononucleated intestines. Because vitellogenins have previously been shown to function redundantly, and downregulation of individual *vit* genes often leads to upregulation of others [26], we analyzed the expression of all six *vit* genes in our dataset and found that all of them were downregulated in worms with a mononucleated intestine (Supplemental Figure 5A-F).

**Figure 4.**
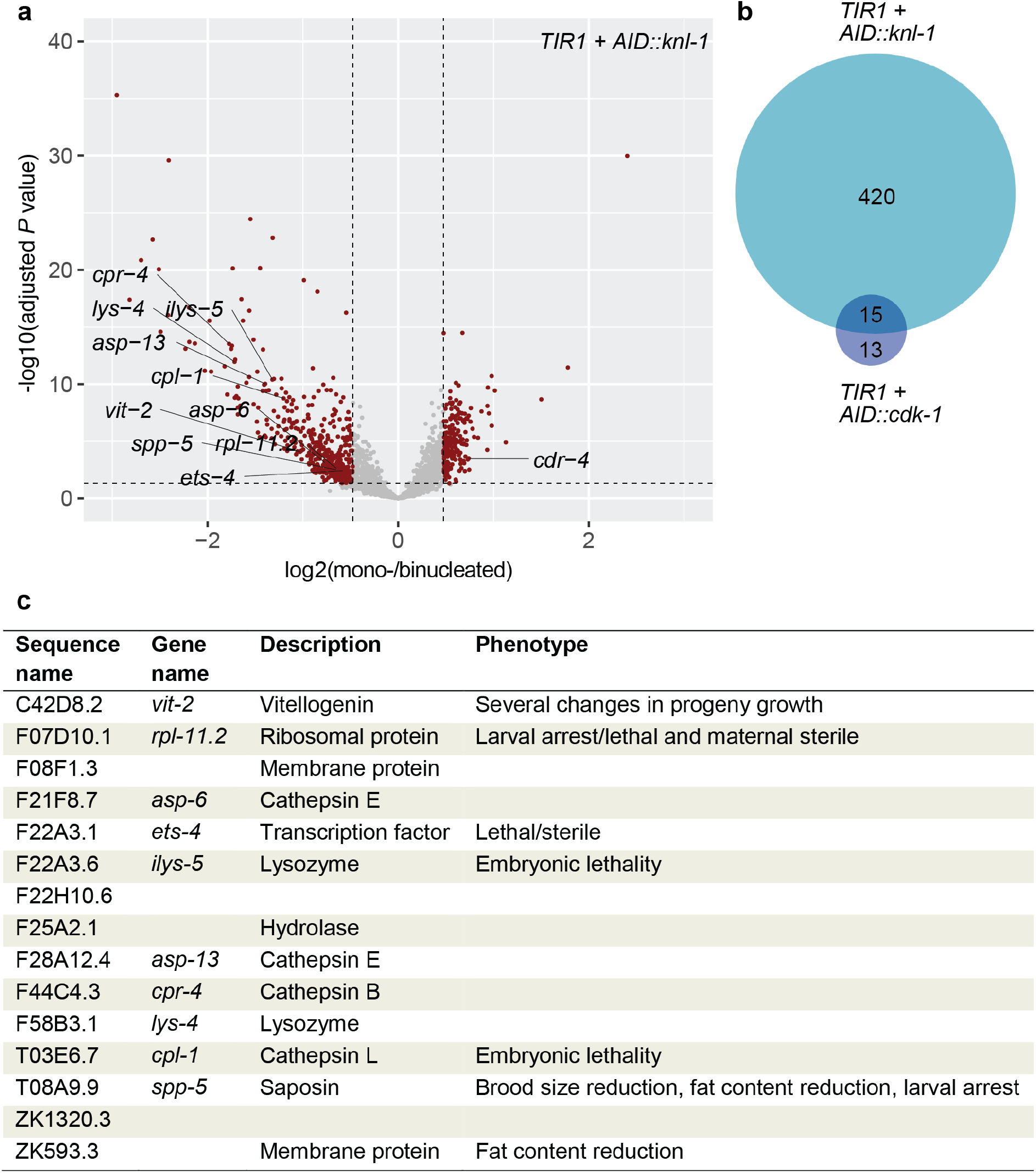
Blocking binucleation causes transcriptional downregulation of intestinally expressed genes in young adult C. elegans. **a.** Volcano plot of RNA sequencing data depicting the transcriptional gene up- and downregulation in worms with a mononucleated intestine, compared to worms with a binucleated (wildtype) intestine in *AID::knl-1 animals* (n = 68). After filtering for coverage, batch consistency and intestinal expression, 2638 genes were analysed for differential gene expression using DEseq. Red dots represent genes significantly differentially expressed (adjusted *P* value <.05, top 25% absolute log2(foldchange)). Genes significantly differentially expressed in both *AID::knl-1* and *AID::cdk-1* comparisons were individually annotated with their gene name (excluding genes without a gene name). **b.** Venn diagram of the overlap of significantly downregulated genes in *AID::knl-1* and *AID::cdk-1* animals. **c.** Overview of genes that are found to be significantly downregulated in worms in which intestinal binucleation is blocked, both by degradation of CDK-1 and degradation of KNL-1, including a short description and previously established associated phenotypes as described on WormBase [30].

Moreover, the sum of all *vit* gene expression values showed a stronger decrease than any gene alone (Supplemental Figure 5G), suggesting that in mononucleated animals, the downregulation of *vit-2* is not compensated by the transcriptional upregulation of other vitellogenins.

### Binucleation of the intestine promotes vitellogenin expression and lipid loading into oocytes

To confirm that *vit-2* expression is downregulated in animals with mononucleated intestinal cells, we generated a *Pvit-2::^NLS^GFP* transcriptional reporter to measure *vit-2* promoter activity at different moments of development. Since vitellogenin expression is absent during larval development and drastically upregulated at the L4-to-adult transition [26], expression levels are still relatively low at 48 hours of development, around the end of the L4 stage (Figure 5A). Upon adulthood, *vit-2* expression levels increase considerably in both auxin-treated and auxin-control animals. However, *vit-2* promoter activity shows a significant reduction in auxin-treated animals at 72 hours of development. Interestingly, the 72-hour timepoint coincides with the moment in development at which worms with a mononucleated intestine produce slow-growing progeny (Figure 3F). Similar to this growth delay, the reduction in vitellogenin expression disappears during the following days of adulthood when *vit-2* expression levels are peaking. Consistent with a transcriptional downregulation of *vit-2* in animals with mononucleated intestinal cells, we also found lower levels of VIT-2 protein in embryos derived from mothers with mononucleated intestines (Figure 5B). Again, VIT-2 levels were significantly lower in embryos derived from auxin-treated 72-hour old adults, but restored to wildtype levels on subsequent days.

**Figure 5.**
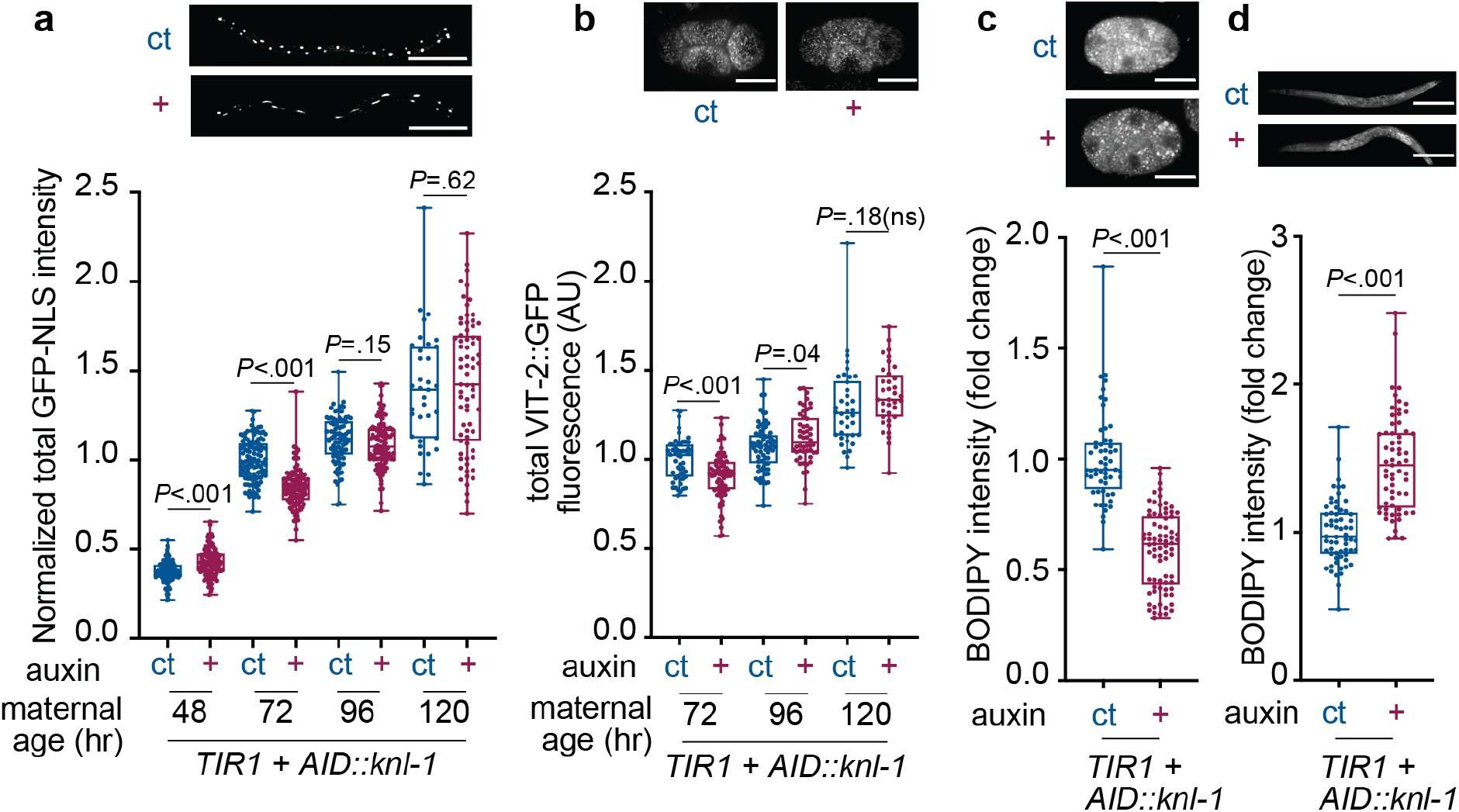
Perturbation of binucleation decreases yolk protein expression and lipid transport to the germline. **a.** Fluorescence images and normalized total fluorescence intensities of a *Pvit-2::GFP* transcriptional reporter at different moments during adult development for worms grown under control (ct, *n* = 53-111) or auxin (+, *n* = 71-111) conditions, in five replicate experiments. Each data point represents the normalized fluorescence intensity of one worm. Boxplots indicate the median and 25^th^-75^th^ percentile, error bars indicate min to max values and individual values are shown as dots. **b.** Fluorescence images and boxplots showing total endogenous VIT-2::GFP fluorescence intensity in early embryos (1-cell stage to 4-cell stage) derived from control (ct, *n* = 38-71) or auxin (+, *n* = 36-71) worms, in two replicate experiments. Boxplots indicate the median and 25^th^-75^th^ percentile, error bars indicate min to max values and individual values are shown as dots. **c.** Z-stack projection of fluorescence images and boxplots of normalized total fluorescence intensities of BODIPY lipid staining of early embryos isolated from young adult (72 hr) worms grown with (+, *n* = 77) or without (ct, *n* = 56) auxin, in three replicate experiments. Boxplots indicate the median and 25^th^-75^th^ percentile, error bars indicate min to max values and individual values are shown as dots. **d.** Z-stack projection images and boxplots of normalized fluorescence intensities of BODIPY lipid staining of young adults (72 hr) that were grown under control (ct, *n* = 62) or auxin (+, *n* = 63) conditions in two replicate experiments. Boxplots indicate the median and 25^th^-75^th^ percentile, error bars indicate min to max values and individual values are shown as dots. *P* values were calculated by unpaired student’s t-test **(a-b)** and Mann-Whitney test **(c-d)**.

Because vitellogenins are required to transport lipids from the intestine to the germline, we investigated whether decreased vitellogenin levels resulted in a reduction of lipid loading into oocytes. For this, we used BODIPY staining to quantify lipid levels in embryos and young adult worms. We observed lower lipid levels in embryos from auxin-treated worms, consistent with reduced lipid loading into oocytes of animals with mononucleated intestines (Figure 5C). Moreover, we found higher amounts of lipids in the intestines of adults with mononucleated intestines (Figure 5D), indicating that specifically the transport of lipids is impaired, rather than lipid production or uptake. The effect of mononucleation on lipid levels was more striking than the decrease that we observed in *Pvit-2::^NLS^GFP* expression, which is consistent with the notion that vitellogenins function in two distinct yolk complexes, and that multiple vitellogenins are downregulated in mononucleated animals. Together, our findings indicate that expression of vitellogenins is hampered by blocking binucleation in the intestine, resulting in reduced lipid loading into developing oocytes.

### Binucleation of the C. elegans intestine is important for rapid upregulation of gene expression

To understand why mononucleated cells have decreased levels of *vit-2* gene expression compared to binucleated cells, we performed in depth analysis of our single-worm RNA sequencing data to identify similarities between genes that are affected by binucleation. First, we found no correlation between expression levels and differential gene expression in animals with a mononucleated intestine, indicating that binucleation is not solely important for the expression of highly or lowly expressed genes (Supplemental Figure 4A,C). Next, we used sequencing data from wildtype worms at different stages of development, available from ModEncode [31], and analyzed the developmental expression profiles of the genes that we identified in our sequencing of animals with mononucleated intestinal nuclei. Overall, genes that are downregulated in worms with a mononucleated intestine show an increase in gene expression throughout wildtype development, with a strong increase in expression from the L4 to adult stage (Figure 6A). In contrast, genes that are not affected by perturbation of binucleation or intestinally expressed genes show a more constant pattern of expression during wildtype development, with intestinal genes even displaying a modest decrease in expression levels throughout the developmental stages. In addition, we found a substantial enrichment (*P*=6.4e-8) of downregulated genes located on the X chromosome (Supplemental Figure 6A). The X chromosome-enriched downregulation is not restricted to vitellogenin genes, of which five out of six are located on the X chromosome, but rather a global repression of all X chromosomal genes (Supplemental Figure 6B). Possibly, an altered X chromosome organization within the nucleus contributes to the decreased expression in mononucleated cells.

**Figure 6.**
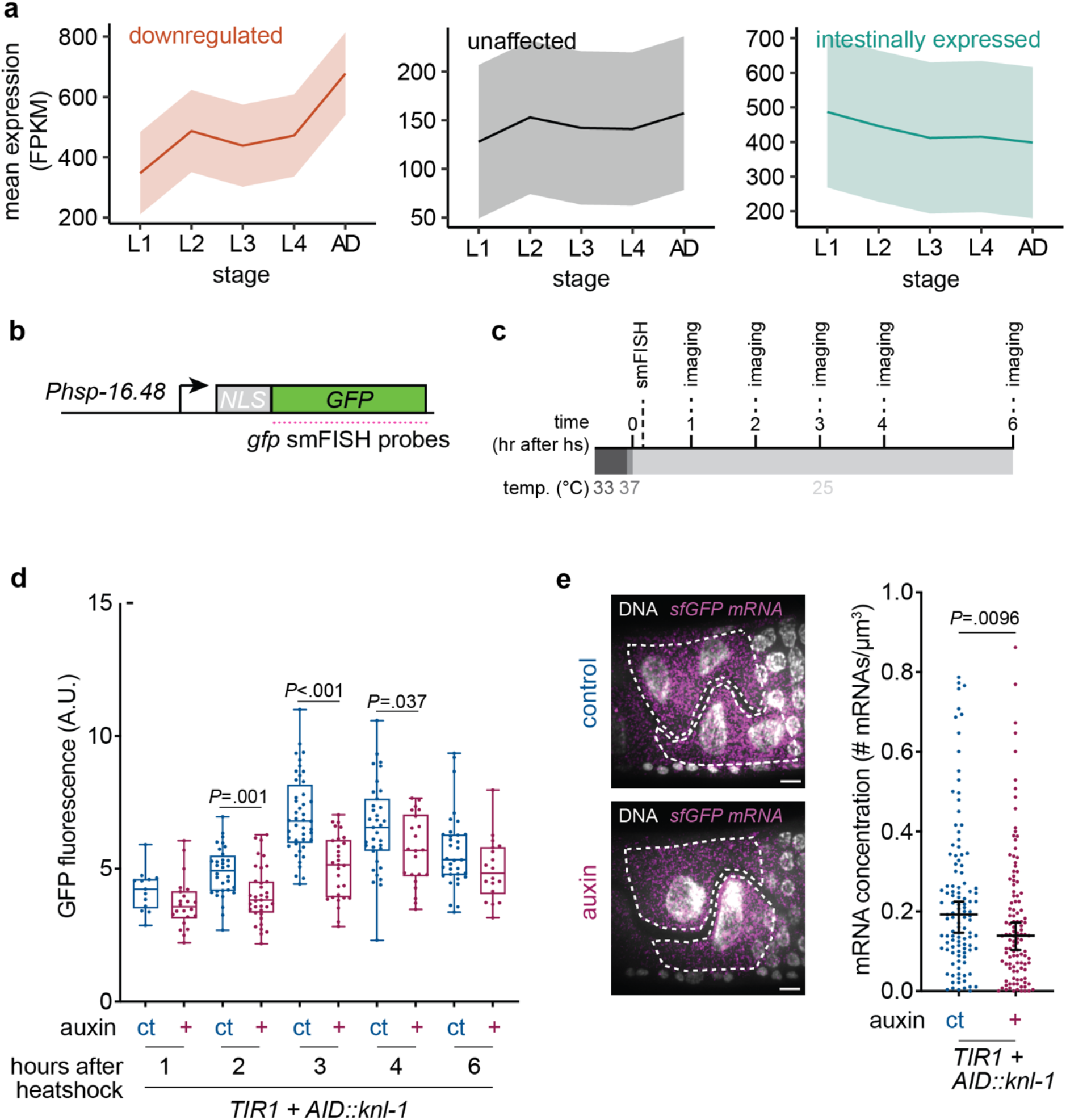
Binucleation allows rapid upregulation of transcription. **a.** Mean expression for downregulated, unaffected or intestinally expressed subsets of genes in different developmental stages from ModEncode FPKM gene expression data. Area around curve shows confidence interval. **b.** Schematic overview of heat shockinducible NLS-GFP transgene. smFISH probes were designed against the sfGFP sequence. **c.** Schematic overview of smFISH and imaging experiments following heat-shock induction. Young L4 animals grown under control or auxin conditions were heatshocked in a waterbath at 33°C for 30 minutes, followed by five minutes in a 37°C air incubator. Animals were left to recover at 25°C. Samples were taken after 15 minutes for smFISH analysis or every hour, for the analysis of nuclear GFP accumulation. **d.** Boxplots of total nuclear fluorescence intensities of intestinal nuclei at different timepoints after heatshock for animals grown under control (ct, *n* = 13-44) or auxin (+, *n* = 18-35) conditions, in three replicate experiments. Boxplots indicate the median and 25^th^-75^th^ percentile, error bars indicate min to max values and individual values are shown as dots. *P* values were calculated by unpaired student’s t-test. **e.** Representative smFISH images and quantifications of cellular *GFP* mRNA concentration in animals grown under auxin (+, *n* = 113) or control (ct, *n* = 112) conditions 15 minutes after heatshock, in two replicate experiments. Scatter plot with line and error bars indicating median and 95% confidence interval, each dot represents a single cell. *P* value was calculated by Mann-Whitney test.

To test whether binucleation is important for rapid upregulation of gene expression in general, or specifically for expression of genes that are upregulated upon adulthood, we measured the upregulation of a heat shock-inducible GFP-NLS reporter in animals with mononucleated or binucleated intestines. In these experiments, L4 stage animals were heat shocked and nuclear GFP intensities were measured at multiple timepoints after heat shock (Figure 6B). Interestingly, mononucleated intestinal cells showed a delay in the upregulation of nuclear GFP after heat shock compared to binucleated controls, whereas the accumulation of GFP signal in body wall muscle nuclei was similar between the two conditions (Figure 6C, Supplemental Figure 8A). To confirm that this delay in upregulation of heat shock-inducible expression arises at the transcriptional level, we quantified the cellular *GFP* mRNA density 15 minutes after heat-shock induction using single molecule fluorescent in situ hybridization (smFISH). Consistent with a delay in GFP upregulation, we observed a significant decrease in *GFP* mRNA levels in mononucleated compared to binucleated intestinal cells (Figure 6D). Together, these results show that rapid upregulation of transcription is impaired in cells with one rather than two nuclei.

A defect in the rapid upregulation of *vit-2* gene expression at the L4-to-adult transition could underlie the reduced vitellogenin expression in mononucleated cells. If so, transient activation of vitellogenin transcription earlier in development should be sufficient to mitigate the differences in *vit-2* expression between binucleated and mononucleated intestines. During wildtype development, vitellogenin expression is upregulated at the L4-to-adult transition by the interaction between transcription factors CEH-60 and UNC-62[32,33]. We found that intestine-specific overexpression of CEH-60 and UNC-62 was sufficient to activate expression of our vitellogenin reporter as early as the L3 stage, whereas no activation was observed at this stage in control animals (Supplemental Figure 9A). Furthermore, animals carrying the *ceh-60;unc-62* overexpression showed an increase in vitellogenin reporter expression levels (Figure 7A), suggesting that these transcription factors are limiting for vitellogenin expression. Importantly, overexpression of CEH-60 and UNC-62 resulted in similar *Pvit-2* expression levels between mononucleated and binucleated intestines, consistent with an impaired activation by these transcription factors in mononucleated cells. Taken together, these results indicate that binucleation of the intestine is needed to facilitate rapid transcriptional upregulation of gene expression at the L4-to-adult transition, which is important for proper intestinal cell function.

**Figure 7.**
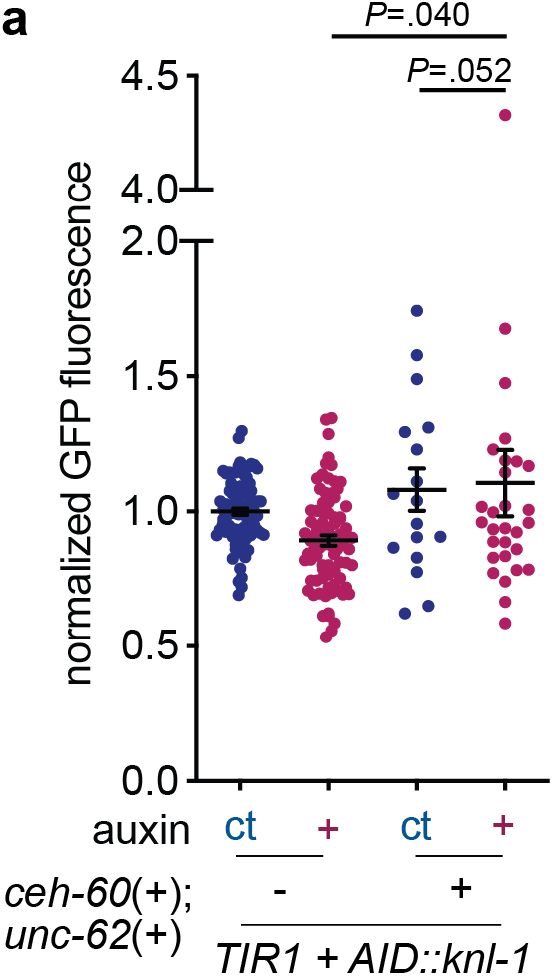
Transient activation of vitellogenin expression rescues vit-2 downregulation in animals with mononucleated intestines. **a.** Normalized total GFP fluorescence intensity per animal, for animals grown under auxin (+) or control (ct) conditions and with (+, *n* = 17 and 29) or without (-, *n* = 82 and 89) an intestinal overexpression of transcription factors *ceh-60* and *unc-62*, measured at young adulthood in five replicate experiments. Scatter plot with line and error bars indicating mean and SEM. *P* values were calculated by Mann-Whitney test.

## Discussion

Our study demonstrates that binucleation is important for rapid transcriptional upregulation of gene expression at the onset of adulthood, indicating that the transcriptional output of a cell is not only determined by the copy number of a gene, but also by whether the DNA is present in a single nucleus or partitioned over two or more nuclei. This finding identifies a key role for endomitosis and multinucleation and highlights the importance of nuclear organization for fine-tuning gene expression control.

Through inhibition of endomitosis in developing *C. elegans* larvae, we were able to compare binucleated and mononucleated intestinal cells with the same genome copy number. Surprisingly, we found that restricting all DNA copies into one nucleus instead of two does not influence cell size or intestinal morphology, but specifically affects the efficiency of fast-acting transcriptional responses. It is possible that differences in nuclear volume or nuclear surface-to-volume cause the transcriptional defects that we observed in mononucleated cells. Changes in the surface-to-volume ratio could influence nuclear import and export dynamics of transcriptional regulators, leading to altered transcriptional dynamics. An increased nuclear volume could lead to lower concentrations of transcription factors in the nucleoplasm, reducing gene target engagement and transcription output. Finally, it has been shown that the expression of developmentally regulated genes often correlates with a specific subnuclear localization and that interactions with the nuclear lamina, the nuclear envelope or the nucleolus can influence transcriptional activity [34–36]. Therefore, changes in the nuclear surface-to-volume ratio and increases in nuclear volume could influence transcription by a differential distribution of loci within the nucleus.

In this study, we show that in *C. elegans* intestinal cells binucleation is important for the rapid upregulation of vitellogenin genes in young adult animals. We find that reduced expression of vitellogenins in mononucleated intestinal cells leads to progeny with developmental delays and reduced fitness. Together, this work sheds light on the function of multinucleation in polyploid cells, revealing that the packaging of nuclear DNA into multiple nuclei can be beneficial to facilitate rapid transcriptional responses. As multinucleated cells are common in many animal and plant tissues, and can also arise in diseases such as cancer, Alzheimer and during viral infections [37–39], our findings raise the possibility that multinucleated tissues function to maximize transcriptional activities in many contexts.

## Online methods

### Worm culture

Worms were cultured on nematode growth medium (NGM) plates seeded with OP50 *Escherichia coli* bacteria according to standard protocols. All strains were maintained at 20°C, unless indicated otherwise. N2 Bristol were used as wildtypes. For auxin plates, NGM was supplemented with 0.5 mM auxin (Sigma-Aldrich I2886) after autoclaving and cooling to <55°C. As auxin is dissolved in 100% ethanol, control plates are supplemented with equal volumes of 100% ethanol without auxin. For auxin experiments, worms were synchronized by isolation of eggs from adult hermaphrodites through alkaline hypochlorite treatment and hatching overnight in M9 medium in the absence of food. Next, synchronized L1 larvae were divided over three possible conditions: no auxin (-), auxin control (ct) and auxin (+). Auxin worms were grown on auxin for 24 hours, after which they are transferred to control plates. Auxin control worms were grown on control plates for 24 hours, and transferred to auxin plates from 24 to 48 hours of development. Worms of the no auxin condition were never grown on auxin plates during their development. Given timepoints can differ based on growth speed of specific transgenic strains. For fitness assays, worms were grown on NGM plates supplemented with 100 mg/mL ampicillin (Sigma-Aldrich A-9518) L4440 *Escheria coli* bacteria to prevent contamination. For heatshock experiments, animals were heatshocked in a waterbath at 33°C for 30 minutes, followed by five minutes in a 37°C air incubator. Animals were then transferred to 25 °C, and samples were taken after 15 minutes for smFISH analysis (see below) or every hour, for the analysis of nuclear GFP accumulation.

### Strains

*C. elegans* strains used in this study are described in Supplementary Table 1. CDK-1 and KNL-1 N-terminal AID tags were inserted into the CA1209 strain using CRISPR/Cas9 genome editing with two overlapping ssODNs (IDT 4 nmole Ultramers) as template for homology-directed repair, as described by Paix and collegues [40]. Single guide RNA and ssODN repair template sequences are listed in Supplementary Table 2. Intestinal markers (*matIs53*), a nuclear membrane marker (*matIs104*), pharyngeal markers (*matIs114* and *matIs116*) or an S/G2-phase marker (*matIs29*) were integrated using gamma-irradiation. A transcriptional *Pvit-2::sfGFP* reporter strain was integrated using CRISPR/Cas9 mediated genome editing at the MosI insertion allele *ttTi5605* on chromosome II.

### Microscopy and image analysis

Static and time-lapse epifluorescence imaging of intestinal fluorescent markers, nuclear markers, chromosome markers, vitellogenin levels and DNA/lipid staining was performed on a PerkinElmer Ultraview Vox spinning disk confocal microscope equipped with a Hamamatsu C13440 digital camera. DIC imaging of worm size was performed using a 20x objective on a Zeiss Axio Imager M2 with Axiocam. Imaging of cell-cycle progression using the S/G2-phase marker was performed on a Leica DM6000 Fluorescence Microscope. For imaging, animals were mounted onto 2% agarose pads (adults) or 7% agarose pads (L1 and L2 larvae) and immobilized with 10 mM sodium azide for static imaging, or 1 ng/mL levamisole and 100 ng/mL tricaine for time-lapse imaging. Young adult intestinal markers or nuclear membrane marker were imaged with a 63x objective, using 2x binning. Chromosome markers in L2/L3 animals were imaged with a 100x objective, without binning. Time-lapse imaging of intestinal markers during endomitosis in L1 larvae was performed with a 100x objective, with 2x binning. For imaging in the presence of auxin, M9 and agarose pads were supplemented with 0.5 mM auxin. To measure vitellogenin and lipid levels, embryos were imaged at 100x magnification, using 2x binning. Propidium iodide DNA staining was imaged with a 63x objective, with 2 x binning. Images were processed and analyzed using Fiji software [41]. For quantifications of PI stainings, GFP-VIT-2 fluorescence intensity, and BODIPY stainings, z-axis serial scans distanced at 50% the optical slice depth were summed in a z-stack projection after background subtraction, after which the integrated density of the region of interest was measured. For quantifications of nuclear GFP after heat-shock induction, 20 z-slices with 0.5 μm spacing (corresponding to 50% of the optical slice depth) were summed in a z-stack projection. In this case, no background subtraction was performed, as background intensity analysis showed similar background levels between conditions. From the z-projections, integrated densities of the nuclear regions were measured. For binucleated cells, the integrated densities of the two nuclei were summed.

### DNA staining

For propidium iodide quantitative DNA staining, freeze-cracked young adult hermaphrodites were fixed with Carnoy’s solution and treated with RNAse A (Invitrogen #60216) as previously described [42]. Propidium iodide staining was performed using 100 μg/mL propidium iodide (Sigma #P4170) for 90 minutes at 37°C, after which slides were washed three times and mounted using ProLong gold antifade mountant (Invitrogen #P36934).

### Competitive fitness assay and modelling

Competitive fitness assays were performed using the *AID::cdk-1* strains GAL182 (with *Pmyo-2::GFP*) and GAL191 (with *Pmyo-2::mCherry*), and the *AID::knl-1* strains GAL160 (with *Pmyo-2::mCherry*) and GAL162 (with *Pmyo-2::GFP*). Prior to the start of the assay, worms were grown under auxin or auxin control conditions as described above. Each experiment contained three conditions: control-GFP versus control-mCherry, auxin-GFP versus control-mCherry and control-GFP versus auxin-mCherry. In the first condition, control-GFP versus control-mCherry, auxin control worms with a green fluorescent pharyngeal marker were grown on the same plate as auxin control worms with a red fluorescent pharyngeal marker. Secondly, for the auxin-GFP versus control-mCherry condition, auxin worms with a green fluorescent pharyngeal marker were grown on the same plate as auxin control worms with a red fluorescent pharyngeal marker. Finally, in the control-GFP versus auxin-mCherry condition, auxin control worms with a green fluorescent pharyngeal marker were grown on the same plate as auxin worms with a red fluorescent pharyngeal marker. For each of the conditions, ten replicates were performed. In each replicate, seven worms of both conditions were transferred to single plates, which were left to grow until starvation (for *AID::knl-1* worms) or until the plate was close to starvation (for *AID::cdk-1* strains, as these were prone to become sterile upon starvation). Each plate was chunked to two new plates, one of which was grown for one day and used to count relative amounts of GFP+ and mCherry+ progeny, whereas the other plate was used to initiate another growth cycle. For counting, animals were washed off plates, mounted on 2% agarose pads containing 10 mM sodium azide, and counted by hand using a Zeiss Axioscope. Relative amounts of control-GFP versus control-mCherry progeny were used to normalize relative amounts of auxin-GFP versus control-mCherry and control-GFP versus auxin-mCherry progeny.

For modelling, the average number of eggs and average time of egg-laying were calculated for the first and second generation, using a predicted gaussian distribution curve on measurements of eggs laid per day. These numbers were used to calculate an exponential growth rate λ, equal to 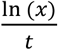, where x is the average number of eggs that one animal produces, and t is the average time (in days) at which these eggs are laid. The predicted relative fitness change per day of growth was determined by the ratio of the growth rate of worms with a mononucleated intestine divided by the growth rate of worms with a binucleated intestine. In the model assuming a transgenerational effect, this change in relative fitness is constant for several generations. When assuming a direct effect, the change in relative fitness is set to zero from the start of generation F2.

### Single worm RNA sequencing and analysis

Transgenic animals were grown until adulthood under auxin or control conditions. Young adult hermaphrodites were picked using tweezers and washed in 0.5 mL sterile demineralized water before adding 200 μL trizol (Invitrogen 15596018). Samples were stored at −80 °C for a limited time. mRNA extraction, barcoding, reverse transcription, *in vitro* transcription and Illumina sequencing library preparation were performed according to the robotized version of the CEL-seq2 protocol [43] using the SuperScript II Double-Stranded cDNA synthesis kit (Thermofisher), Agencourt AMPure XP beads (Beckman Coulter) and randomhexRt for converting aRNA to cDNA using random priming. The libraries were sequenced paired-end at 50 bp read length on an Illumina HiSeq 2500. The 50 base pair paired-end reads were aligned to the *C. elegans* reference transcriptome compiled from the *C. elegans* reference genome WS249. Raw data was processed using R (v3.6.2). Samples were filtered to include those with sufficient counts per transcript and gene, results were filtered for batch consistency and only intestinally-expressed genes that were found at least once in every sample were included for downstream analysis (Supplemental Figure 3). Additionally, possibly growth-delayed or sterile outlier animals were removed from the dataset, based on spermatogenesis and oogenesis-related gene expression [44]. This resulted in a dataset consisting of 143 animals with gene expression data for a total of 2638 genes for *AID::knl-1* animals and 2724 genes for *AID::cdk-1* animals. The DEseq R package was used to perform differential gene expression analysis. GO term enrichment analysis was performed using the InterMineR package for R. FPKM gene expression data from different developmental stages was retrieved from ModEncode through WormBase and analyzed in R. A Venn diagram was constructed using the BioVenn web application [45].

### Lipid staining

Embryo lipid staining was performed as described previously [25]. In short, embryos were obtained by alkaline hypochlorite treatment of adult hermaphrodites, transferred to PCR tubes (Eppendorf) containing 4% formaldehyde in M9 buffer and subjected to three freeze-thaw cycles in liquid nitrogen and a 37 °C water bath. Embryos were then incubated with 1 μg/ml BODIPY 493/503 (Thermo Fisher Scientific) in M9 buffer for one hour and washed three times with M9 + 0.01% Triton X-100 (brand). Next, nuclei were stained with 5 ng/mL DAPI (brand) and imaged by epifluorescence microscopy.

Adults were fixed for lipid staining by incubating in 4% paraformaldehyde (Sigma-Aldrich 158127) for 1 hour at room temperature. Adults were then washed twice with PBS, resuspended in 95% ethanol and washed again in PBS. Next, fixed animals were incubated with 1 μg/ml BODIPY 493/503 (Thermo Fisher Scientific) in M9 buffer for one hour and washed three times with M9 + 0.01% Triton X-100 (Sigma #9001-93-1). Animals were stained with 5 ng/mL DAPI (brand) and imaged by epifluorescence microscopy.

### Viability assay

Worms were grown until the L4 stage under auxin, control or no auxin condition as described above. Single hermaphrodites were then transferred to new plates every 24 hours. On each day, the number of live progeny and unhatched eggs were counted 24-48 hours after removal of the adult worm to calculate the total brood size and embryonic lethality.

### Progeny growth assay

Animals were grown until young adults under auxin, control or no auxin condition as described above. Developmental transition timings were performed as previously described using a wash-off staging [25]. Briefly, plates containing synchronized gravid hermaphrodites were washed to remove all animals except embryos that remained attached to the solid media. To tightly synchronize a population of animals, larvae that had hatched within one hour after wash-off were collected and transferred to new plates. The fraction of animals that had undergone the transition from L3 to L4 was counted for three replicate plates containing around 150 worms each, in three separate experiments. To estimate the time at which half of the population had undergone the developmental transition, each population was scored at least twice, once before and once after half of the population had undergone the L4 transition. Count data was modelled using a binomial generalized linear model, and the time at which half of the population had undergone the developmental transition (T50) was estimated using the function ‘dose.p’ of the library ‘MASS’ in R. 95% confidence intervals were estimated by multiplying the standard error obtained by the ‘dose.p’ function by 1.96. The significance of the difference between two calculated T50 values was determined using the ‘comped’ function in the library ‘drc’ in R, which takes as input the two T50 values, their standard error and a desired probability level (such as 0.95).

### Single molecule fluorescence in situ hybridization (smFISH)

smFISH was performed as previously described with some minor adjustments. In short, animals were fixed using 4% paraformaldehyde, resuspended in 70% ethanol and stored at 4°C for up to two weeks. Short oligonucleotide probes complementary to the sfGFP sequence (see Supplemental Table 3 for sequences) were designed using a web-based algorithm (www.biosearchtech.com/stellaris-designer) and ordered with a 3-amino modification to enable coupling to Cy5 (GE Healthcare, cat. no. PA25001) as previously described [46]. Hybridization was performed overnight at 37°C in the dark, after which samples were washed and stained with DAPI. Z-stacks of Int3A and Int3V cells were generated using spinning disk microscopy and mRNA molecules were counted using the batch analysis mode in FISH-quant [47]. Before analysis, 4.5 μm substacks were made in Fiji of each cell centered around the nucleus/nuclei. From these substacks, mRNA spots were counted in three regions of 4.55 μm x 4.55 μm, and spot density was calculated as # of spots per μm^3^.

### Statistics and reproducibility

The sample size (*n*) as well as the number of replicate experiments performed for each experiment is described in the corresponding figure legend. All statistical analyses were performed using GraphPad Prism (8.4.3) except for sequencing data analyses. Quantitative data displayed as boxplots indicate the median and 25^th^-75^th^ percentile, error bars indicate min to max values and individual values are shown as dots, unless indicated otherwise. Twotailed unpaired Student’s t-tests were used for pairwise comparisons between groups with similar Gaussian distributions. Mann-Whitney U tests were used for pairwise comparisons between groups with non-Gaussian distributions. The type of statistical test performed is indicated in the corresponding figure legends. Statistical significance values (*P*) are shown directly within the figure above pairwise comparisons.

## Acknowledgements

We thank members of the Galli, Korswagen and Kops labs for helpful discussions, Tim Hoek for help and feedback on fitness assay modelling, and Marvin Tanenbaum for critical reading of the manuscript. We also thank Anko de Graaff and the Hubrecht Imaging Center (HIC) for assistance with microscopy. The spinning disk microscope is funded by equipment grant 834.11.002 from the Dutch Organization of Scientific Research (NWO). We thank the Temmerman lab for sharing the plasmid containing *Pelt-2::ceh-60*. Some strains were provided by the CGC, which is funded by the NIH Office of Research Infrastructure Programs (P40 OD010440). This work was supported by funding from the Human Frontiers Science Program Organization to M.G. (CDA00018/2017-C), an NWO-Veni grant to M.G. (016.Veni.181.016) and a ZonMW grant to M.G. (ETH 40-43500-98-4087).

