## Supplemental Figures and Tables for "Endomitosis controls tissue-specific gene expression during development"

### Supplemental information

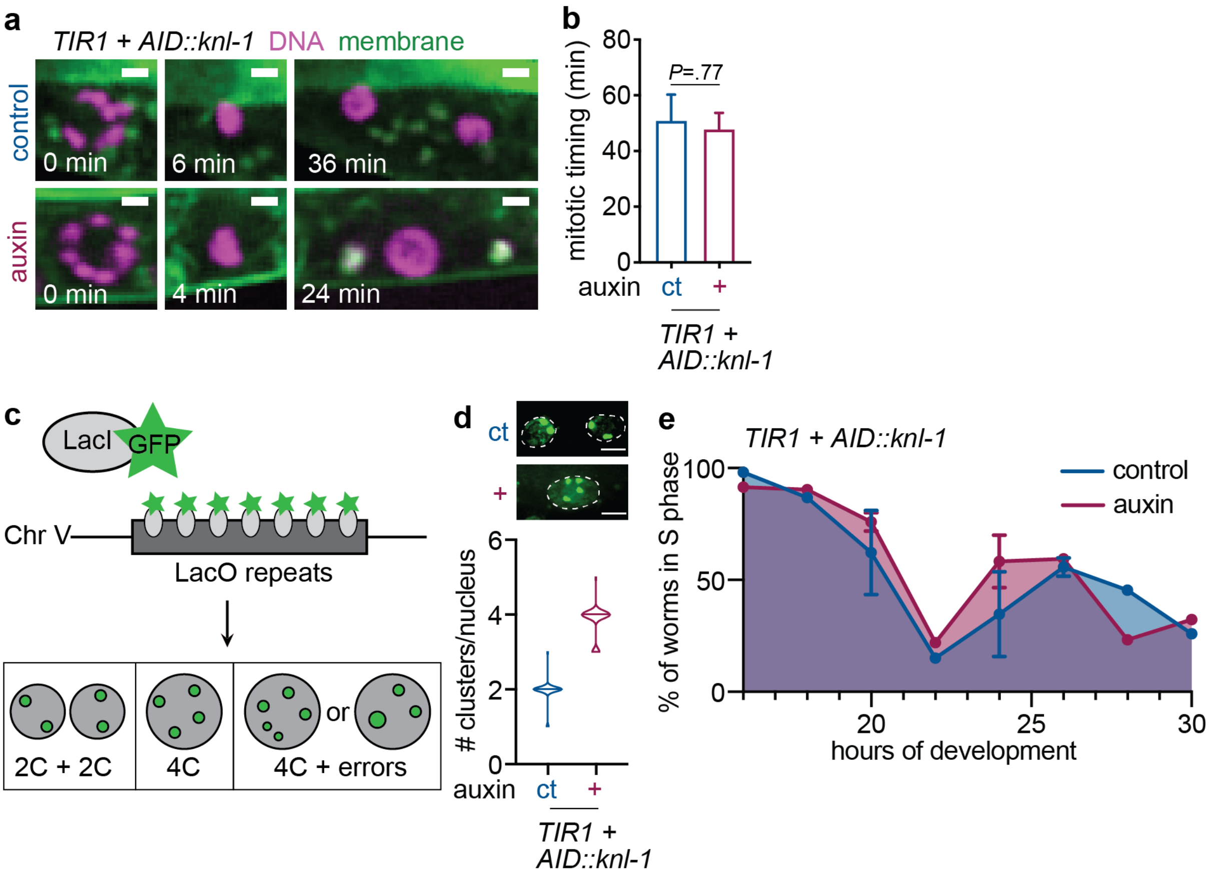


**Supplemental Figure 1.** *Perturbation of binucleation through degradation of KNL-1 prevents DNA segregation and does not affect subsequent S-phase timing.* **a.** Stills of time-lapse videos of cells undergoing endomitosis in the absence (control) or presence of auxin, showing intestinal H2B-mCherry (DNA, shown in magenta) and GFP-PH (membrane, shown in green). Scale bar is 2 µm. Time stamp indicates time after nuclear envelope breakdown. **b.** Quantification of mitotic timing, the duration from nuclear envelope breakdown to nuclear envelope reformation, in the absence (ct, $n = 4$) or presence (+, $n = 7$) of auxin. Bar graph showing mean, error bars indicate SEM. $P$ values were calculated by unpaired student's t-test. **c.** Overview of LacI/LacO system for detection of individual chromosomes in polyploid cells. A series of LacO repeats present on chromosome V are visualized upon heat-shock induced expression of a LacI fused to GFP. After endomitosis in L1, a binucleated cell with two 2C nuclei or a mononucleated cell with a single 4C nucleus should show four individual chromosomes if no segregation errors occurred. If errors did occur during segregation, an alternate number of individual chromosomes should be visible. **d.** Fluorescent images and violin plots showing chromosome cluster counts of fluorescent LacI::GFP foci in control (ct, $n = 98$) and auxin-treated (+, $n = 82$) worms. To distinguish LacI::GFP signal from cytoplasmic autofluorescent vesicles, only nuclear dots were counted as chromosome clusters. Error bars represent min and max values, horizontal bars represent median. Scale bar is 5 µm. **e.** Average percentage of animals in which intestinal cells are undergoing G2 or S phase, determined by the presence of CYB-$1^{DB}$::mCherry, during endomitotic and subsequent endoreplicative cycles in the first and second larval stage for 60 to 200 worms per condition per timepoint, in three replicate experiments. X axis starts at 16 hours. Error bars represent standard deviation.

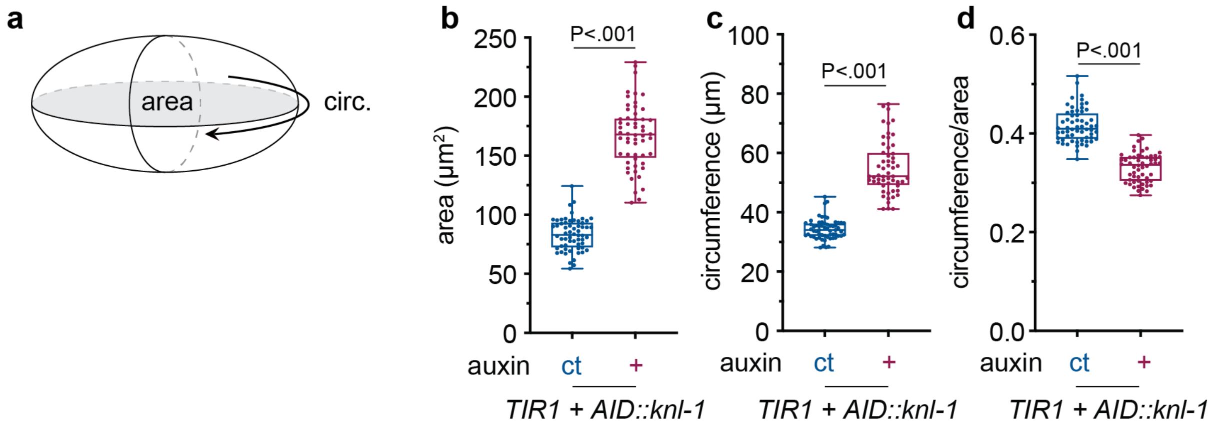


**Supplemental Figure 2.** *Mononucleation results in an altered circumference-to-area ratio.* **a.** Schematic depicting the nuclear parameters measured, area and circumference of a nuclear midplane sections. **b-d.** Boxplots depicting nuclear section area **(b)**, circumference **(c)** and nuclear circumference-to-area ratio **(d)** in binucleated (ct, $n$ = 60) or mononucleated (+, $n$ = 56) cells. Measurements were made at the midplane of the nucleus. Boxplots indicate the median and 25$^{th}$-75$^{th}$ percentile, error bars indicate min to max values and individual values are shown as dots. $P$ values were calculated by Mann-Whitney **(b-c)** and unpaired student's t-test **(d)**.

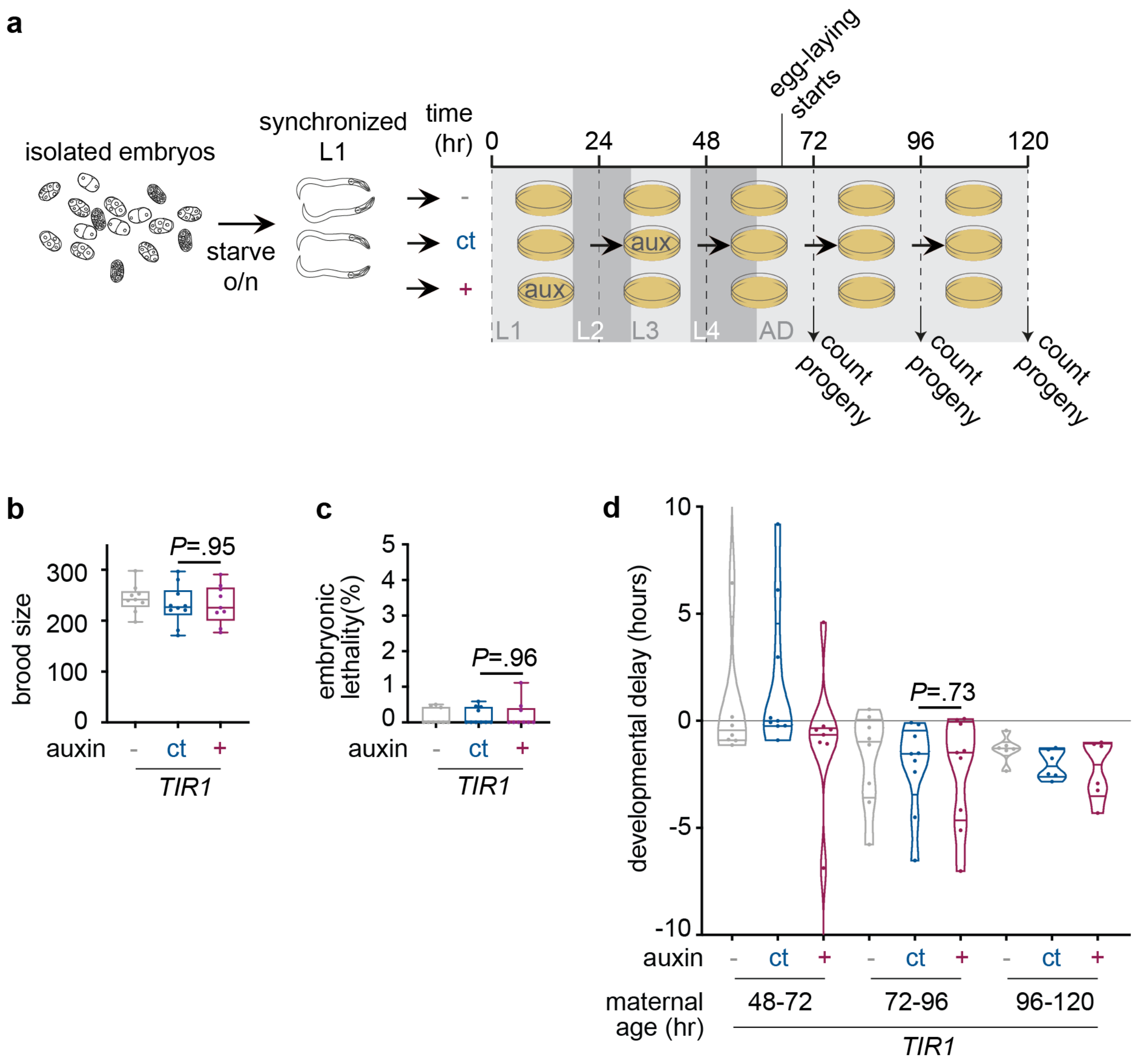


**Supplemental Figure 3.** *Growing worms on auxin in the absence of an AID tag does not affect reproduction or progeny growth.* **a.** Overview of experimental procedures for quantifications of brood size and embryonic lethality. Isolated embryos are synchronized as L1's and divided over the three conditions. Worms start laying eggs between 48 and 72 hours of development. Worms are transferred to new plates at 72, 96 and 120 hours and progeny and eggs on the previous plate is counted after an incubation time of 16-18 hours. **b-c.** Boxplots showing total brood size **(b)** and embryonic lethality **(c)** of worms carrying an intestinally expressed TIR1, grown in the absence of auxin (-, $n$ = 9 plates), under control conditions (ct, $n$ = 10 plates) or in the presence of auxin (+, $n$ = 9 plates). Amounts of eggs and hatched animals were counted for three days of egg-laying. Boxplots indicate the median and 25$^{th}$-75$^{th}$ percentile, error bars indicate min to max values and individual values are shown as dots. $P$ values were calculated by Mann-Whitney test. **d.** Violin box plots depicting progeny growth rates of animals derived from mothers of different ages, which were grown without auxin (-, $n$ = 18 plates), under control conditions (ct, $n$ = 18 plates) or in the presence of auxin (ct, $n$ =18 plates), in three replicate experiments. Horizontal lines indicate the median and 25$^{th}$-75$^{th}$ percentile, violin plots extend to min and max values and individual values are shown as dots. $P$ value was calculated by Mann-Whitney test.

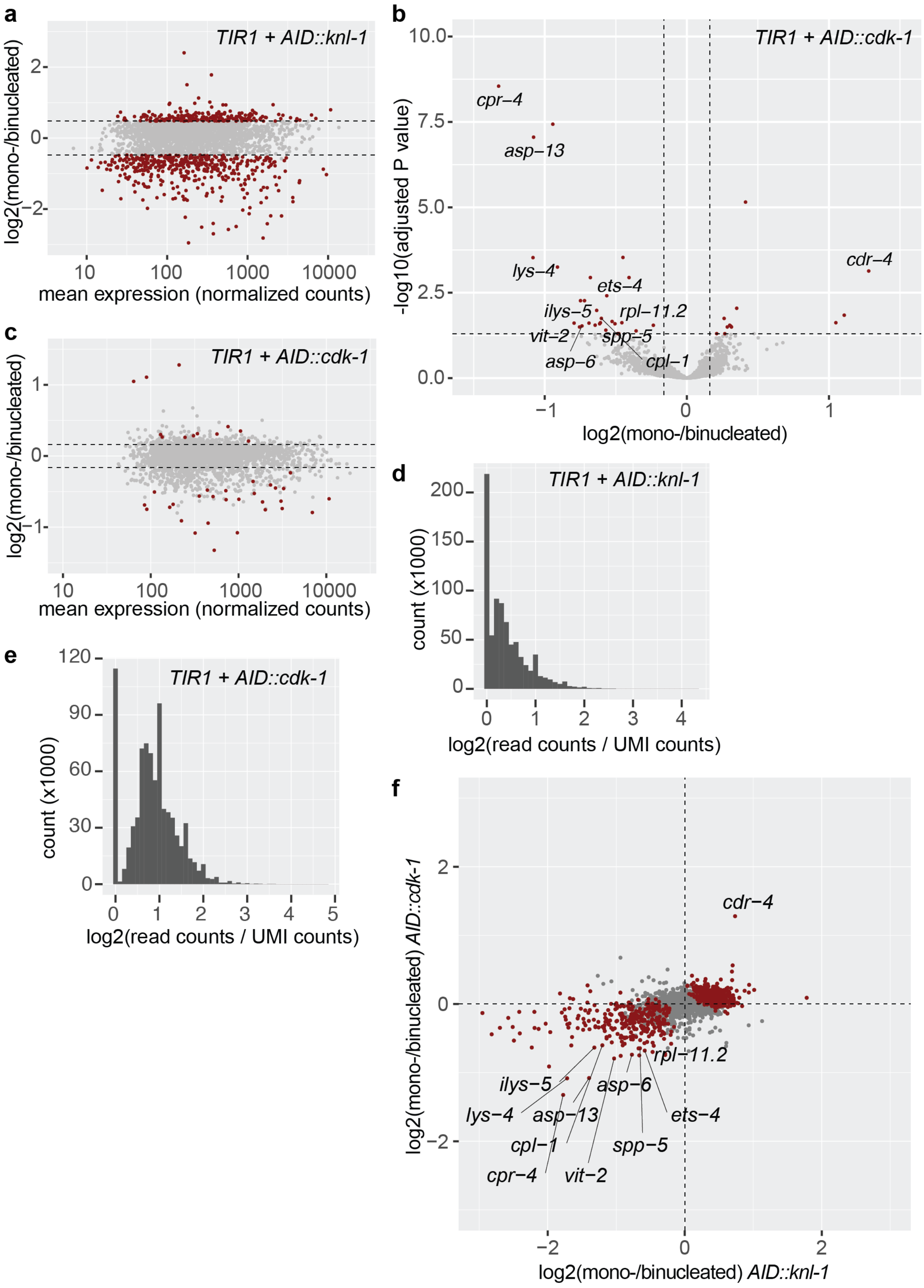


**Supplemental Figure 4.** *Overlap in differential gene expression of single young adult worms containing either AID::knl-1 or AID::cdk-1.* **a-c.** Volcano and MA plots of RNA sequencing data depicting the transcriptional gene up- and downregulation in worms with a mononucleated intestine, compared to worms with a binucleated (wildtype) intestine in animals containing either *AID::knl-1* (**a**) or *AID::cdk-1* (**b-c**), in relation to gene expression levels. Red dots represent genes that are differentially expressed with an adjusted *P* value below .05 and belong to the top 25% regarding absolute log2(foldchange). **d-e.** Histogram depicting the log2 of read counts per unique molecular identifier (UMI) for each gene found in animals containing either *AID::knl-1* (**d**) or *AID::cdk-1* (**e**). **f.** Dot plot depicting the correlation between differential expression in *AID::knl-1* (x-axis) and *AID::cdk-1* (y-axis) animals with a mononucleated versus binucleated intestine. Red dots represent genes that are differentially expressed in the combined dataset with an adjusted *P* value below .05. Genes significantly differentially expressed in both *AID::knl-1* and *AID::cdk-1* comparisons individually were annotated with their gene name (excluding genes without a gene name).

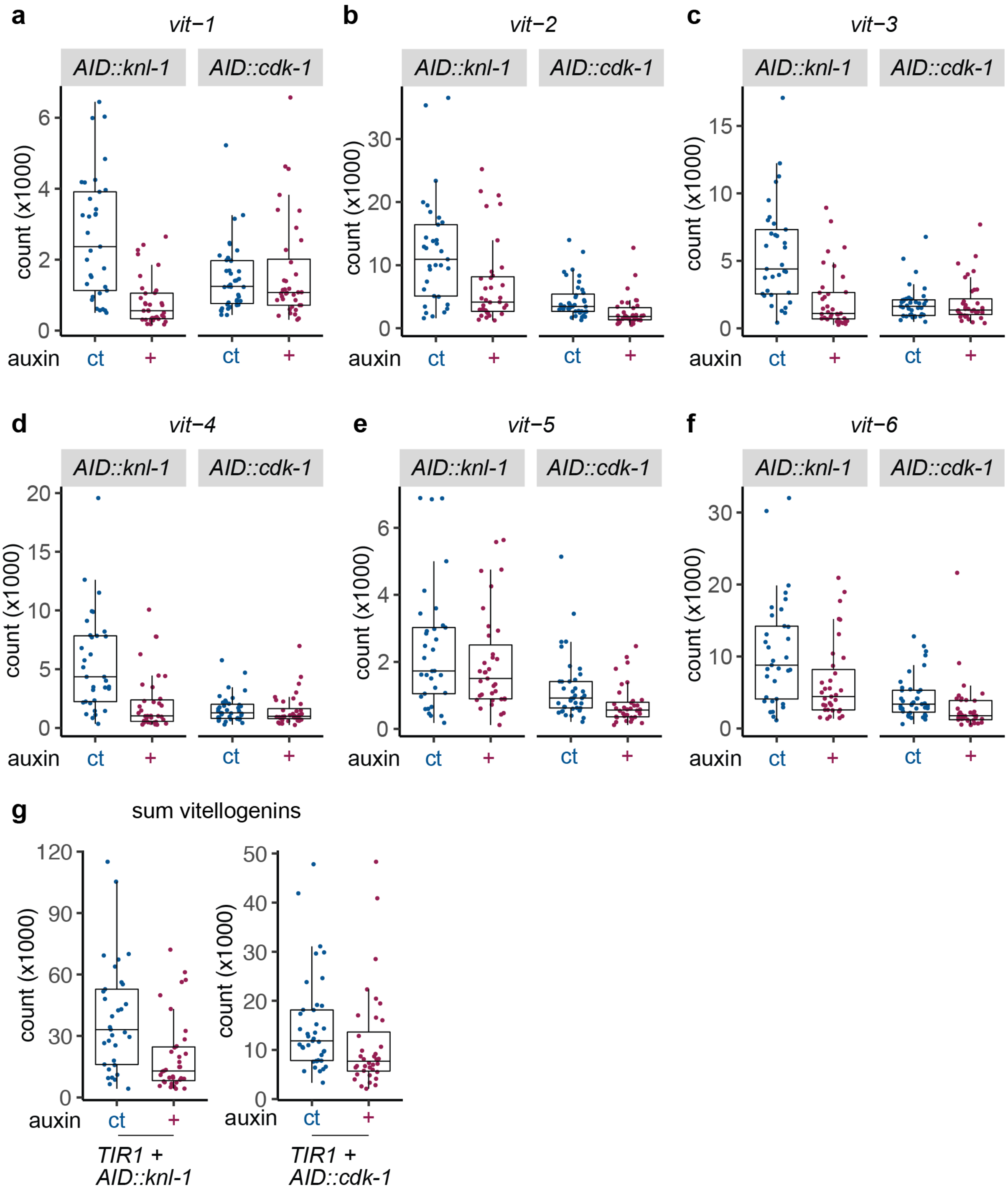


**Supplemental Figure 5.** *Differential expression of vitellogenin genes in RNA sequencing of animals with a mononucleated versus binucleated intestine.* **a-g.** Tukey boxplots showing the separate **(a-f)** and accumulated **(g)** RNA expression levels for all six vitellogenin genes (*vit-1* through *vit-6*) in auxin-control (ct, *n* = 33 for *AID::knl-1* and n=39 for *AID::cdk-1*) or auxin-treated (+, *n* = 35 for *AID::knl-1* and n = 36 for *AID::cdk-1* ). Each dot represents the expression in one worm.

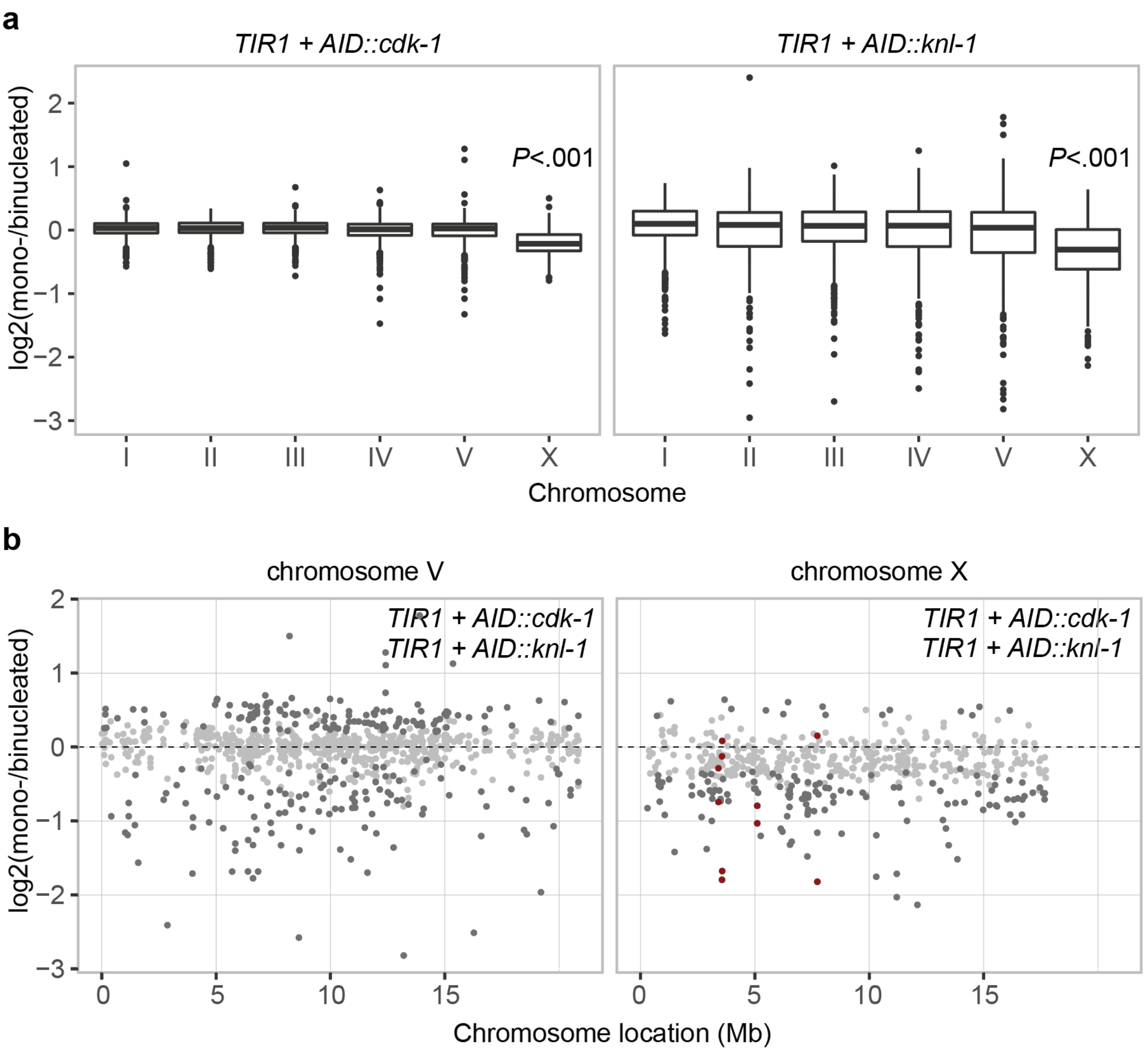


**Supplemental Figure 6.** *Mononucleation of the intestine causes an X chromosome specific decrease in expression levels.* **a.** Tukey boxplots showing the log2(foldchange) differential expression of genes in animals with a mononucleated versus worms with a binucleated intestine per chromosome in *AID::cdk-1* and *AID::knl-1* strains. *P* values of the comparison between X chromosomal and autosomal differential gene expression were calculated by Wilcoxon rank sum test. **b.** Compiled log2(foldchange) of differential expression of genes located on chromosome V and X in animals with a mononucleated versus binucleated intestine. Data is compiled from single worm RNA sequencing data of both *AID::cdk-1* and *AID::knl-1* strains. Genes with a top 25% percent absolute log2(foldchange) are indicated in dark grey, vitellogenins are indicated as red dots.

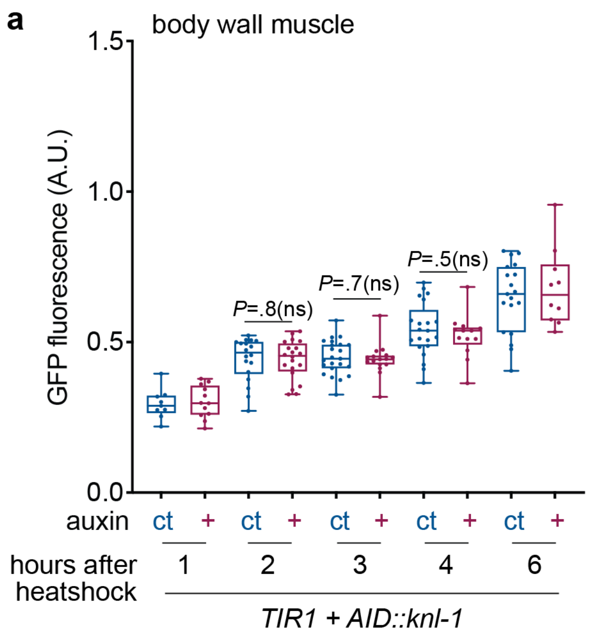


**Supplemental Figure 7.** *Heat-shock response in body wall muscle cells is not affected by mononucleation of the intestine.* **a.** Boxplots with total nuclear fluorescence intensities of body wall muscle nuclei from at different timepoints after heatshock in auxin-control (ct, *n* = 9 - 23) or auxin-treated (+, *n* = 10-20) animals, performed in three replicate experiments. Boxplots indicate the median and 25th-75th percentile, error bars indicate min to max values and individual values are shown as dots. *P* values were calculated by Mann-Whitney test.

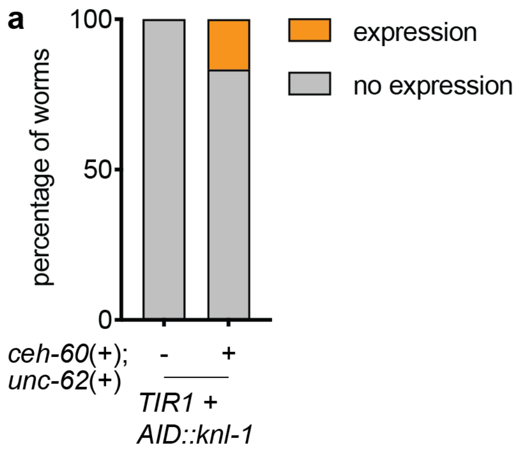


**Supplemental Figure 8.** *Early expression of ceh-60 and unc-62 induces vitellogenin promoter activity before adulthood.* **a.** Bar graph depicting the percentage of animals with high levels of normalized total GFP fluorescence intensity per animal containing *Pvit-2::GFP-NLS*, for L3 animals with (+, *n* = 36) or without (-, *n* = 165) an intestinal overexpression of transcription factors *ceh-60* and *unc-62*. A high level of total GFP fluorescence intensity is defined as more than twice the average control levels of GFP fluorescence intensity after background subtraction.

Supplementary Table 1.

| Strain name | Genotype | Source/ reference |
|---|---|---|
| BCN9071 | *vit-2(crg9070[vit-2::gfp]) X.* | *Caenorhabditis* Genetics Center [25] |
| CA1209 | *ieSi61 [Pges-1::TIR1::mRuby::unc-54 3'UTR + Cbr-unc-119(+)] II; unc-119(ed3) III* | *Caenorhabditis* Genetics Center [16] |
| GAL71 | *ieSi61 [Pges-1::TIR1::mRuby::unc-54 3'UTR + Cbr-unc-119(+)] II; cdk-1(hu262 [AID::cdk-1]) III* | This study |
| GAL117 | *knl-1(mat91[AID::KNL-1]) III; ieSi61 [Pges-1::TIR1::mRuby::unc-54 3'UTR + Cbr-unc-119(+)] II; unc-119(ed3) III* | This study |
| GAL126 | *knl-1(mat91[AID::knl-1]) III; ieSi61 [Pges-1::TIR1::mRuby::unc-54 3'UTR + Cbr-unc-119(+)] II; unc-119(ed3) III; matIs53[Pges-1::sfGFP-PH::tbb-2 3'UTR; Pges-1::H2B-mCherry::unc-54 3'UTR; Plin-48::TdTom]] X* | This study |
| GAL137 | *knl-1(mat91[AID::KNL-1]) III; ieSi61 [Pges-1::TIR1::mRuby::unc-54 3'UTR + Cbr-unc-119(+)] II; unc-119(ed3) III; vit-2(crg9070[vit-2::gfp]) X* | This study |
| GAL141 | *matIs105 [Pges-1::Tir1-tagBFP]; matIs104 [Pact-5::npp-9-mCherry::tbb-2 3'UTR] II; knl-1(mat91[AID::KNL-1]) III* | This study |
| GAL160 | *matIs114 [Pmyo-2::mCherry]; ieSi61 [Pges-1::TIR1::mRuby::unc-54 3'UTR + Cbr-unc-119(+)] II; knl-1(mat91[AID::KNL-1]) III* | This study |
| GAL162 | *matIs116 [Pmyo-2::GFP]; ieSi61 [Pges-1::TIR1::mRuby::unc-54 3'UTR + Cbr-unc-119(+)] II; knl-1(mat91[AID::KNL-1]) III* | This study |
| GAL163 | *knl-1(mat91[AID::KNL-1]) III; ieSi61 [Pges-1::TIR1::mRuby::unc-54 3'UTR + Cbr-unc-119(+)] II; matIs29 [Pges-1::CYB-1 DB::mCherry::unc-54 3' UTR; Pges-1::NLS-sfGFP::tbb-2 3' UTR; Pmyo-2::GFP]* | This study |
| GAL178 | *knl-1(mat91[AID::KNL-1]) III; ieSi61 [Pges-1::TIR1::mRuby::unc-54 3'UTR + Cbr-unc-119(+)] II; unc-119(ed3) III; syIs44 [Phsp-16::lacI::GFP + lacO + dpy-20(+)] V* | This study |
| GAL182 | *matIs115 [Pmyo-2::GFP]; ieSi61 [Pges-1::TIR1::mRuby::unc-54 3'UTR + Cbr-unc-119(+)] II; cdk-1(hu262 [AID::cdk-1]) III* | This study |
| GAL191 | *matIs114 [Pmyo-2::mCherry]; ieSi61 [Pges-1::TIR1::mRuby::unc-54 3'UTR + Cbr-unc-119(+)] II; cdk-1(hu262 [AID::cdk-1]) III* | This study |
| GAL192 | *matIs137[Pvit-2::NLS-sfGFP-AID_P2A_NLS-sfGFP-AID::tbb-2 3'UTR] II; ieSi61 [Pges-1::TIR1::mRuby::unc-54 3'UTR + Cbr-unc-119(+)] II; knl-1(mat91[AID::knl-1])III* | This study |

| | | |
|---|---|---|
| GAL225 | *matIs155[Phsp-16.48::NLS-sfGFP::tbb-2 3' UTR]; knl-1(mat91[AID::knl-1])III; ieSi61 [Pges-1::TIR1::mRuby::unc-54 3'UTR + Cbr-unc-119(+)] II; unc-119(ed3) III* | This study |
| GAL226 | *matIs137[Pvit-2::NLS-sfGFP-AID_P2A_NLS-sfGFP-AID::tbb-2 3'UTR] II; ieSi61 [Pges-1::TIR1::mRuby::unc-54 3'UTR + Cbr-unc-119(+)] II; knl-1(mat91)[AID::KNL-1] III; matEx151 [Pelt-2::ceh-60; Pges-1::BFP-P2A-unc-62; Pmyo-2::mCherry]* | This study |
| TY5434 | *syIs44 [Phsp-16::lacI::GFP + lacO + dpy-20(+)] V.* | *Caenorhabditis* Genetics Center [48] |

Supplementary Table 2.

| Description | Sequence |
|---|---|
| *cdk-1* N-terminal guide RNA target sequence | ataggatccataactaaaat |
| *cdk-1* repair ssODN 1 | ccttttcggcggcagtggaaatgattcaaaattacacgcttacgccttttctatcgttgtattacaattgttctgacaaaattcatttccAattttagttATGCCTAAAGATCCAGCCAAACCTCCGGCCAAGGCACAAGTTGTGGGATGGCCACCGGTGAGATCATACCGGAAGAACGTGATGG |
| *cdk-1* repair ssODN 2 | GCCACCGGTGAGATCATACCGGAAGAACGTGATGGTTTCCTGCCAAAAATCAAGCGGTGGCCCGGAGGCGGCGGCGTTCGTGAAGGATCCTATTCGCGAAGGAGAAGTGGCCCACGAGGGAGATTCGGTTTACACACTCAACGATTTCACGAAGCTCGAAAAAATCGGCGAAGGAACATACGGAG |
| *knl-1* N-terminal guide RNA target sequence | cttacgaggctccatcgaca |
| *knl-1* repair ssODN 1 | tttattaccatttttaaaacatatttacagccatgCCTAAAGATCCAGCCAAACCTCCGGCCAAGGCACAAGTTGTGGGATGGCCACCGGTGAGATCATACCGGAAGAACGTGATGGTTTCCTGCC |
| *knl-1* repair ssODN 2 | GAGATCATACCGGAAGAACGTGATGGTTTCCTGCCAAAAATCAAGCGGTGGCCCGGAGGCGGCGGCGTTCGTGAAGggaggagccggagcatcgatggagcctcgtaagaagcggaactcgattct |

Supplementary Table 3

| smFISH probe sequences for *sfGFP* (28 probes) | | | |
|---|---|---|---|
| tggattagcttttcgacgtc | gcaagctttttagcattttc | caaccttgcgtttctttttc | agaattgggacaactccagt |
| cccattaacatcaccatcta | cctctccacggacagaaaat | taagggtgagttttccgttt | gtagttttccagtagtgcaa |
| agcattgaacaccataggtc | tcatgtgatccggataacgg | gggcatggcactcttgaaaa | agtgcgttcctgtacataac |
| tcccgtcatctttgaaagat | aaacttgacttcagcacgcg | gattaacaagggtatcacct | tcaataccctttaactcgat |
| agaatgtttccatcttcttt | agttgtactcgagtttgtgt | gccgtgatgtatacattgtg | gattccattcttttgtttgt |
| acggaaccatcttcaacgtt | gttgataatggtctgctagt | ccatcgccaattggagtatt | ggttgtctggtaaaaggaca |
| acagattgtgtcgacaggta | ttttcgttgggatctttcga | tcaagaaggaccatgtggtc | aatcccagcagcagttacaa |
